# PHLPP2 is a pseudophosphatase that lost activity in the metazoan ancestor

**DOI:** 10.1101/2024.12.03.625870

**Authors:** Tarik Husremović, Katharina M. Siess, Sumire Antonioli, Vanessa Meier, Lucas Piëch, Irina Grishkovskaya, Nikoleta Kircheva, Silvia E. Angelova, Andreas Brandstätter, Jiri Veis, Fran Miočić-Stošić, Dorothea Anrather, Markus Hartl, Linda Truebestein, Bojan Žagrović, Stephan Hann, Christoph Bock, Egon Ogris, Todor Dudev, Nicholas A.T. Irwin, David Haselbach, Thomas A. Leonard

## Abstract

The phosphoinositide 3-kinase (PI3K) pathway is a major regulator of cell and organismal growth. Consequently, hyperactivation of PI3K and its downstream effector kinase, Akt, is observed in many human cancers. PH domain leucine-rich repeat-containing protein phosphatases (PHLPP), two paralogous members of the metal-dependent protein phosphatase family, have been reported as negative regulators of Akt signaling and, therefore, tumor suppressors. However, the stoichiometry and identity of the bound metal ion(s), mechanism of action, and enzymatic specificity of these proteins are not known. Seeking to fill these gaps in our understanding of PHLPP biology, we unexpectedly discovered that PHLPP2 has no catalytic activity against the regulatory phosphorylation sites of Akt, nor the generic substrate *para*-nitrophenylphosphate. Instead, we found that PHLPP2 is a pseudophosphatase with a single zinc ion bound in its catalytic center. Furthermore, we found that current cancer genomics data do not support the proposed role of PHLPP1 or PHLPP2 as tumor suppressors. Phylogenetic analyses revealed an ancestral phosphatase that arose more than 1000 Mya, but that lost activity at the base of the metazoan lineage. In summary, our results provide a molecular explanation for the inconclusive results that have hampered research on PHLPP and argue for a new focus on non-catalytic roles of PHLPP1 and PHLPP2.

**Significance Statement:** PHLPP1 and PHLPP2 have previously been reported as protein phosphatases that specifically inactivate Akt, a pro-growth and survival kinase hyperactivated in many human cancers. Unexpectedly, we found that purified PHLPP2 has no detectable enzymatic activity in vitro, an observation which can be rationalized by its unusual active site, which has diverged significantly from that of canonical metal-dependent phosphatases. Furthermore, we show that cancer genomics do not support a role for either PHLPP1 or PHLPP2 in cancer. Our findings argue for the exploration of alternative hypotheses regarding the role of PHLPP in Akt signaling and cancer, with a focus on its non-catalytic functions.

## Introduction

Cell and organismal growth depend on growth factors that activate receptor tyrosine kinases (RTKs) expressed on the surface of cells. RTK activation leads to the recruitment and activation of the small GTPase Ras and the lipid kinase phosphoinositide 3-kinase (PI3K), which synergistically drive the conversion of phosphatidylinositol-4,5- bisphosphate (PIP_2_) in the plasma membrane into the lipid second messenger phosphatidylinositol-3,4,5-trisphosphate (PIP_3_). PIP_3_, in turn, recruits and activates a number of effector proteins, including the serine/threonine protein kinase Akt and its upstream activator phosphoinositide-dependent kinase 1 (PDK1) (1). PDK1 and Akt control essential aspects of cell growth, proliferation, differentiation and metabolism, including glucose homeostasis (2). The lipid phosphatase and tumor suppressor, phosphatase and tensin homolog (PTEN), attenuates PI3K signaling by converting PIP_3_ back into PIP_2_. Mutations in Ras and PI3K, as well as loss of PTEN are frequently observed in human cancers (3). Akt itself is also found mutated in cancer (4) and the rare overgrowth disorder, Proteus Syndrome (5).

Akt is an AGC kinase, consisting of a PIP_3_-binding pleckstrin homology (PH) domain and a kinase domain which bears two regulatory phosphorylation motifs. PDK1 phosphorylates Akt1 at T308 in its activation loop (6), while mTORC2 is widely believed to phosphorylate S473 in its hydrophobic motif (7). A third phosphorylation site in the C-terminal tail, the turn motif, is constitutively phosphorylated and controls Akt stability (8). Phosphorylation of T308 and S473 promote disorder-to-order transitions of their respective motifs, leading to Akt activation (9).

PIP_3_ turnover by PTEN leads to the dissociation and inactivation of Akt. An important, but poorly understood aspect of Akt inactivation is the dephosphorylation of T308 and S473 by protein phosphatases. T308 is dephosphorylated by protein phosphatase 2A (PP2A) (10, 11), whereas S473 has been reported to be the target of the PH domain leucine-rich repeat-containing protein phosphatases PHLPP1 and PHLPP2 (12, 13). The original identification of PHLPP as an Akt phosphatase was guided by the hypothesis that, since S473 phosphorylation is growth factor or serum-sensitive, such a phosphatase might contain a PH domain. The PH domain would presumably target it to the same membranes where Akt is activated (12). While PHLPP1 and PHLPP2 double knockout mice exhibit only a mild colitis resistance phenotype (14), their proposed repressive effects on Akt signaling have led to their designation as tumor suppressor genes (12, 15, 16). However, the mechanisms by which these phosphatases target Akt for dephosphorylation and the structural basis for their specificity are currently unknown.

Here, we show that – in contrast to previous reports – PHLPP2 is actually a pseudophosphatase. We show that PHLPP2 has no activity in vitro against either phosphorylated T308 or S473 of Akt1, and that cancer genomics do not support a role for either PHLPP1 or PHLPP2 in cancer, which questions their designation as tumor suppressors in the PI3K/Akt pathway. We corroborate the absence of phosphatase activity by showing that PHLPP evolved from an ancestral phosphatase that lost activity in the metazoan ancestor. In summary, our results refute the notion that PHLPP is an Akt phosphatase, question efforts to pharmacologically target PHLPP activity (15, 17), and establish a molecular and evolutionary basis for reassessing the biological roles of PHLPP1 and PHLPP2 as pseudophosphatases.

## Results

### PHLPP2 exhibits no detectable activity against Akt

We expressed GST-tagged, full-length human PHLPP2 in baculovirus-infected Sf9 insect cells and purified it to homogeneity by sequential affinity, ion exchange and size-exclusion chromatography (Supplementary Figure 1A). PHLPP2 was confirmed to be predominantly monomeric by mass photometry (Supplementary Figure 1B) and the masses of all recombinant PHLPP2 constructs were verified by denaturing mass spectrometry (Supplementary Figure 1C-E). PHLPP2 and all mutants were confirmed to be properly folded by analytical size exclusion chromatography (Supplementary Figure 1F) and thermal stability assays (Supplementary Figure 1G). Addition of EDTA to PHLPP2 did not affect its global thermal stability (Supplementary Figure 1G).

Akt has been reported as a substrate of both PHLPP1 and PHLPP2 in mammalian cells (12, 13, 18). Figure 1A shows the structure of the kinase domain of Akt alongside the sequence motifs of its activation loop and hydrophobic motif. According to reports, PHLPP is specific for the hydrophobic motif (pS473). To test the activity of recombinant PHLPP2 against synthetic Akt phosphopeptides, we monitored phosphate production in a malachite green assay. As a control, we assayed a mutant of PHLPP2 (D1024N) (Supplementary Figures 1E), designed to abrogate catalytic activity (18). Unexpectedly, PHLPP2^D1024N^ exhibited comparable activity to wild-type PHLPP2 against the Akt1 activation loop peptide (Figure 1B). When assayed against both the activation loop and hydrophobic motif peptides, we detected very low, but variable activity from multiple independent preparations of wild-type PHLPP2. This activity, however, was entirely lost upon treatment of each preparation with 12.5 nM okadaic acid (Figure 1C-D), a highly specific PP2A/PP4/PP6 inhibitor and toxin produced by several species of dinoflagellate (19, 20). Previous reports that PP2A dephosphorylates T308 of Akt1 (10, 11, 21) are consistent with both the higher activity against the activation loop peptide and inhibition by okadaic acid. As a control, we assayed the activity of recombinant PP2A against both peptides, which exhibited a specific activity 50-fold greater than any of our PHLPP2 preparations (Figure 1C-F). Inhibition of all phosphatase activity in our PHLPP2 assays occurred despite PHLPP2 being present in molar excess over okadaic acid in all but one concentration. Finally, PHLPP2 exhibited no activity against either pT308 or pS473 of stoichiometrically phosphorylated full-length Akt1 [prepared according to ref (22)] under the assay conditions of the study that reported PHLPP phosphatase activity for the first time (12) (Supplementary Figure 2A-C).

**Figure 1.**
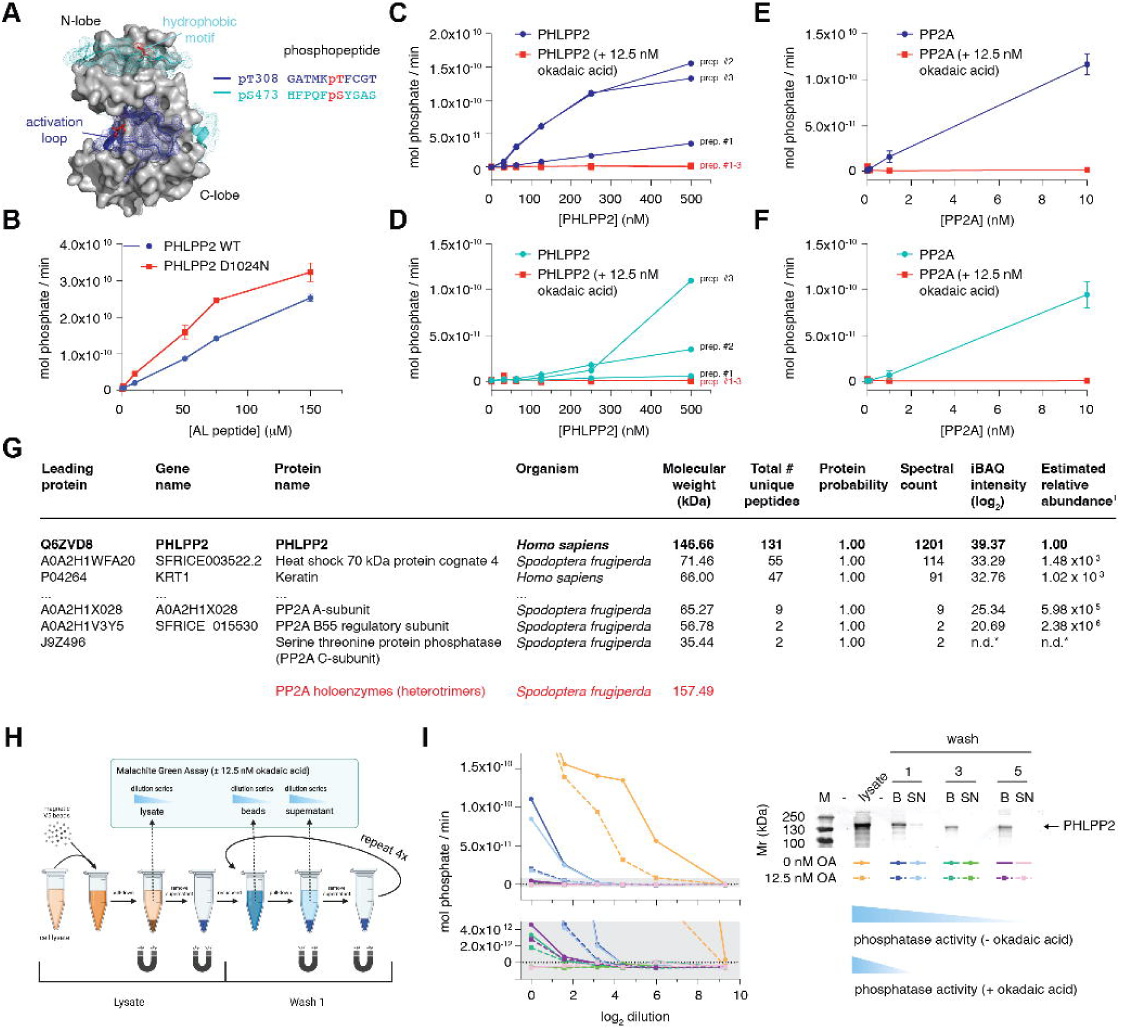
PHLPP2 exhibits no detectable activity against Akt. A. Structure of Akt1 kinase domain depicting activation loop (T308, blue) and hydrophobic motif (S473, cyan) phospho-peptides used in phosphatase assays. B. Protein phosphatase activity of purified PHLPP2^D1024N^ (red) compared to PHLPP2^WT^ (blue) against the activation loop peptide of Akt. C. Dephosphorylation of the activation loop peptide by three independent preparations of recombinant PHLPP2 purified from Sf9 insect cells (blue) ± 12.5 nM okadaic acid (red). D. Dephosphorylation of the activation loop peptide by purified PP2A (blue) ± 12.5 nM okadaic acid (red). E. Dephosphorylation of the hydrophobic motif peptide by three independent preparations of recombinant PHLPP2 purified from Sf9 insect cells (cyan) ± 12.5 nM okadaic acid (red). F. Dephosphorylation of the hydrophobic motif peptide by purified PP2A (cyan) ± 12.5 nM okadaic acid (red). G. Peptide fingerprinting mass spectrometry to identify proteins in recombinant PHLPP2 purified from insect cells. Red: molecular weight of Sf9 PP2A heterotrimers. H. Schematic for immunoprecipitation of PHLPP2 from HEK293 cells and subsequent phosphatase assay. I. Phosphatase activity of immunoprecipitated PHLPP2 determined after successive wash steps ± 12.5 nM okadaic acid. B = beads, SN = supernatant.

Suspecting that recombinant PHLPP2 preparations may be contaminated with one or more phosphatases of the PP2A family, we subjected purified PHLPP2 to peptide fingerprinting mass spectrometry. While PHLPP2 was estimated to be more than 99.9% pure, peptides of all three subunits of *Spodoptera frugiperda* PP2A were detected (Figure 1G). A total of 13 unique peptides corresponding to the PP2A catalytic (C) subunit (2), A-subunit (9) and B55 (B) regulatory subunit (2) were measured. With a mass of 157 kDa, the PP2A heterotrimer is very close in mass to PHLPP2 (147 kDa), such that they cannot be separated by size-exclusion chromatography. The identification of PP2A holoenzymes in recombinant PHLPP2 preparations explains the okadaic acid sensitivity of the observed phosphatase activity (Figure 1C-D).

To assess whether contaminating phosphatase activity may have been the source of literature reports of PHLPP activity, we immunoprecipitated ectopically expressed V5-tagged PHLPP2 from HEK293 cells using V5 nanobody-conjugated magnetic beads and subjected the beads to a series of successive wash steps, while measuring phosphatase activity at each step in the absence and presence of okadaic acid (Figure 1H). We observed that all of the measurable phosphatase could be successively washed away and that any residual activity could be inhibited by okadaic acid (Figure 1I; Supplementary Figure 2D). In summary, we conclude that high purity preparations of recombinant PHLPP2 as well as PHLPP2 immunoprecipitated from mammalian cells can easily be contaminated with PP2A.

### PHLPP2 is a pseudophosphatase

To confirm that the activity detectable in our purified PHLPP2 came from contaminating PP2A, we modified our purification protocol to include the reverse affinity-based removal of PP2A. Produced by the cyanobacterium, *Microcystis aeruginosa,* microcystin-LR is a high-affinity irreversible inhibitor of PP1, PP2A, PP3, PP4 and PP6 phosphatases that forms a covalent bond to PP2A C269 (Figure 2A) and binds to PP2A in the same binding site as okadaic acid (Figure 2B) (20). Microcystin-LR can be covalently coupled to sepharose beads to create phosphatase inhibitor beads (PIBs, Figure 2C) (23). We confirmed that our PIBs were highly effective in enriching PP2A from PHLPP2-expressing Sf9 cell lysates by peptide fingerprinting mass spectrometry (Figure 2D). As a control we treated an equivalent lysate with 500 nM okadaic acid to inhibit all okadaic acid-sensitive phosphatases. Since okadaic acid and microcystin-LR bind to the same pocket, PIBs should not enrich these phosphatases in the okadaic acid-treated control. Mass spectrometry analysis confirmed a 1200-, 541- and 65-fold enrichment of the PP2A C, A and B subunits respectively. The three most enriched proteins correspond to the same three contaminating PP2A proteins observed in PHLPP2 preparations (Figure 2D, Supplementary Figure 2E). Importantly, recombinant PHLPP2 was not observed to be enriched over the control lysate (Figure 2D).

**Figure 2.**
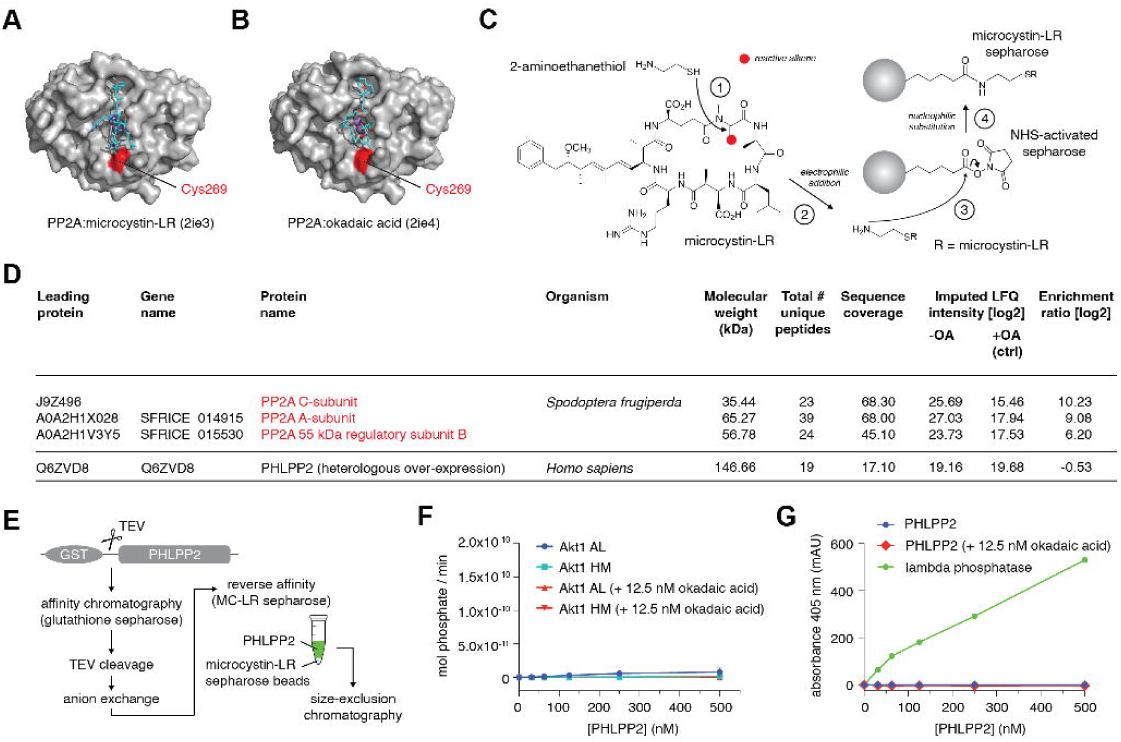
PHLPP2 is a pseudophosphatase. A. Structure of PP2A catalytic subunit bound to the covalent inhibitor microcystin-LR. B. Structure of PP2A catalytic subunit bound to okadaic acid. C. Schematic for covalently coupling microcystin-LR to NHS-activated sepharose beads (23). D. Tandem mass spectrometry analysis of proteins bound to microcystin-LR- conjugated beads after incubation with cell lysates of Sf9 cells heterologously over-expressing PHLPP2. E. Purification scheme for affinity-based removal of PP2A family phosphatases from purified PHLPP2. F. Dephosphorylation of activation loop (blue) and hydrophobic motif (cyan) peptides by PHLPP2 purified according to the scheme in K ± 12.5 nM okadaic acid (red). G. Dephosphorylation of the generic phosphatase substrate para-nitrophenyl phosphate (pNPP, blue) by PHLPP2 purified according to the scheme in K ± 12.5 nM okadaic acid (red) or lambda phosphatase (green). Assay buffer includes 2 mM MnCl_2_.

To remove contaminating PP2A from PHLPP2, we incubated the purified protein with PIBs following anion exchange chromatography and prior to size-exclusion chromatography (Figure 2E). PIB-treated PHLPP2 lost all detectable activity against both Akt phosphopeptides (Figure 2F), even at concentrations 500-fold above its known physiological concentration (24). Finally, we evaluated the capacity of PHLPP2 to dephosphorylate *para*-nitrophenyl phosphate (pNPP). PHLPP2 that had been purified via the scheme shown in Figure 2E exhibited no activity against pNPP even in the absence of okadaic acid (Figure 2G). As a positive control, pNPP was robustly dephosphorylated by lambda phosphatase under the same conditions (Figure 2G). Importantly, since lambda phosphatase depends on manganese ions for its activity (25), which is also the case for PP2C family phosphatases (26, 27), this experiment explicitly rules out that PHLPP2 exhibits activity in the presence of millimolar concentrations of manganese. In summary, we were unable to detect any phosphatase activity associated with purified, PP2A-free PHLPP2.

### PHLPP2 is a zinc-binding protein with altered metal ion stoichiometry

Members of the metal-dependent family of protein phosphatases (PPM), exemplified by protein phosphatase Mg^2+^/Mn^2+^-dependent (PPM1A), depend on two or three divalent metal ions in their active site for catalysis (26–28) (Figure 3A). PHLPP1 and PHLPP2, however, exhibit poor conservation of a number of key residues required for metal ion coordination (Figure 3B).

**Figure 3.**
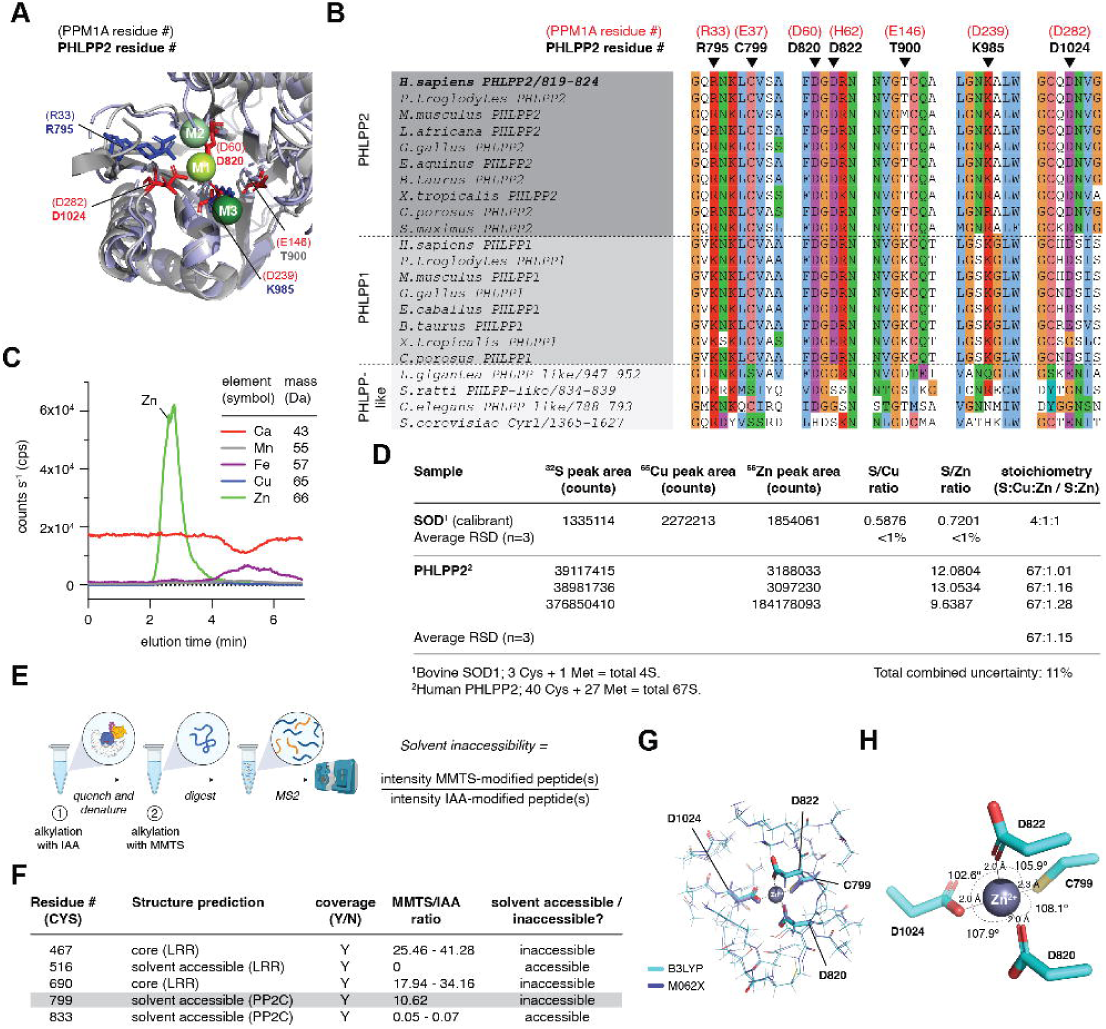
PHLPP2 is a zinc-binding protein. A. Catalytic site of PPM1A phosphatase (grey) superimposed on AF2 model of PHLPP2 PP2C domain (lilac). M1, M2 and M3 metal ions shown in spheres; metal coordinating amino acids shown in stick representation. PPM1A residue numbers shown in parentheses. B. Multiple sequence alignment of the metal ion-coordinating motifs of PHLPP1, PHLPP2, and PHLPP-like proteins across a wide evolutionary time. PPM1A residue numbers shown above the alignment in parentheses. C. ICP-MS chromatograms for recombinant human PHLPP2. D. Calibration of PHLPP2:Zn^2+^ stoichiometry for three biological replicates. E. Differential alkylation of PHLPP2 under native and denaturing conditions, coupled to mass spectrometry. F. Differential alkylation of five different peptides of PHLPP2 (two representing buried cysteines, 2 representing surface exposed cysteines, and one encompassing C799). G. Quantum mechanical simulation of zinc ion coordination sphere using the ONIOM method (30). Results of two quantum mechanical frameworks (B3LYP and M062X) are shown. H. Geometry of the zinc-binding site from the simulations showing bond angles and lengths.

To investigate whether the metal ion-binding properties of PHLPP2 could explain the lack of catalytic activity, we determined the identity and stoichiometry of any metal ions co-purifying with PHLPP2 by size-exclusion chromatography coupled to inductively-coupled plasma mass spectrometry (SEC-ICP-MS). This analysis revealed a single zinc ion in three independent preparations of the protein (Figure 3C-D; Supplementary Figure 3A-B). No signal for iron, copper, manganese, or calcium was detected (Figure 3C).

Using the AlphaFold2-predicted structure of the phosphatase domain, which superimposes on the experimentally determined structure of PPM1A with a root mean square deviation (r.m.s.d.) of 1.09 Å, the most likely constellation of zinc-coordinating residues was identified as C799, D820, D822 and D1024, which correspond to the M2 metal ion binding site (Supplementary Figure 3C). The crystal structure of the related pseudophosphatase, [Transforming growth factor β-activated kinase 1]-binding protein 1 (TAB1), also revealed a single manganese ion in the M2 site (29) (Supplementary Figure 3D). The M1 and M3 binding sites can be excluded on the basis of a lack of conservation of suitable residues at these positions. All four putative Zn^2+^-coordinating residues (C799, D820, D822, and D1024) are invariant in chordate PHLPP1 and PHLPP2 genes (Figure 3B). No other candidate cysteine, histidine, aspartate or glutamate residues exist in the vicinity of the M2 site.

To determine experimentally whether C799 was a coordinating ligand of the zinc ion, we alkylated purified PHLPP2 under native conditions with iodoacetamide (IAA), followed by methyl methanethiosulfonate (MMTS) under denaturing conditions (Figure 3E) and analyzed the peptides by tandem mass spectrometry (Figure 3F, Supplementary Figure 3E). C467 and C690 exhibited very high ratios of MMTS- to IAA- modified peptides, consistent with them being buried in the hydrophobic core of the LRR domain. Conversely, residues in the intrinsically disordered C-terminal tail all exhibited very low ratios, consistent with high solvent accessibility. C799, though predicted to be solvent exposed, exhibited a high MMTS to IAA ratio, indicating that it is protected from alkylation under native conditions. We therefore concluded that C799, in the M2 binding site, coordinates the single zinc ion that co-purifies with PHLPP2.

To obtain a reasonable starting structure for the bound zinc ion, we performed explicit-solvent molecular dynamics (MD) simulations starting from the predicted PHLPP2 structure and using distance restraints between the ion and the coordinating atoms obtained from the structure of the C1D3 zinc finger of α-klotho (30). During the simulations, the global atom-positional root-mean-square deviation (RMSD) with respect to the initially equilibrated structure reached a stable level of 0.2-0.25 nm after 10 ns, indicating that placement of the zinc ion in the M2 site is compatible with the folded conformation of the PP2C domain (Supplementary Figure 3F). Since MD is not capable of modeling the electronic environment with sufficient accuracy, we modeled the binding site using the ONIOM approach (31). Here, atoms three bonds away from the zinc ion were treated quantum-mechanically, while atoms further away were treated semi-empirically. To validate the ONIOM method, we simulated the zinc binding site of α-klotho (30). The best scoring models of the zinc binding site obtained from two quantum mechanical frameworks exhibited near tetrahedral geometry and bond lengths consistent with experimentally determined zinc clusters (Figure 3G-H, Supplementary Figure 3G-H). The free energy change associated with substitution of zinc for either manganese (Mn^2+^) or iron (Fe^2+^) was 7-12 kcal/mol (Supplementary Figure 3I), indicating that the binding site is energetically optimized for zinc.

Since all known PPM family phosphatases require a minimum of two metal ions (M1 and M2) for catalytic activity, the absence of PHLPP2 activity can be explained.

### Cancer genomics does not support a role for PHLPP1 or PHLPP2 as tumor suppressors

Since we could not measure any detectable phosphatase activity of PHLPP2, we investigated the evidence for the role of PHLPP1 or PHLPP2 as tumor suppressors. To assess somatic mutations in either PHLPP1 or PHLPP2 associated with cancer, we plotted synonymous (silent) and non-synonymous (missense) mutations reported in the Catalog of Somatic Mutations In Cancer (COSMIC) database (32) and compared them to two Cancer Gene Census (CGC) (33) tumor suppressor genes, two CGC oncogenes, and two randomly chosen olfactory receptor genes as negative controls (genes that are very unlikely to be functionally involved in cancer development). Figure 4A shows that the rate of both synonymous and non-synonymous mutations in *PHLPP1* and *PHLPP2* is comparable to those of the olfactory receptor genes *OR2A4* and *OR51E2*, while the rates of non-synonymous mutations in both *PTEN* and *TP53*, as well as *KRAS* and *BRAF*, are significantly higher in human cancers. As expected, non-synonymous loss-of-function (LOF) mutations are significantly enriched at hotspots (red) known to disrupt the function of *PTEN* and *TP53* tumor suppressor genes, while gain-of-function (GOF) mutations are enriched in *KRAS* and *BRAF* oncogenes (Figure 4B). In contrast, *PHLPP1* and *PHLPP2* show comparable rates of non-synonymous mutations to *OR2A4* and *OR51E2* throughout their coding regions. To determine whether copy number variants (CNVs) are significantly enriched in human cancers, we plotted copy number losses (CNL) and gains (CNG) for each of the genes in Figure 4A. *PTEN* and *TP53* exhibit a high frequency of CNL (red) consistent with their tumor suppressive function, while *KRAS* and *BRAF* exhibit a high frequency of CNG (green) consistent with their oncogenic potential (Figure 4C). In contrast, *PHLPP1* and *PHLPP2* show comparable rates of CNVs to *OR2A4* and *OR51E2*.

**Figure 4.**
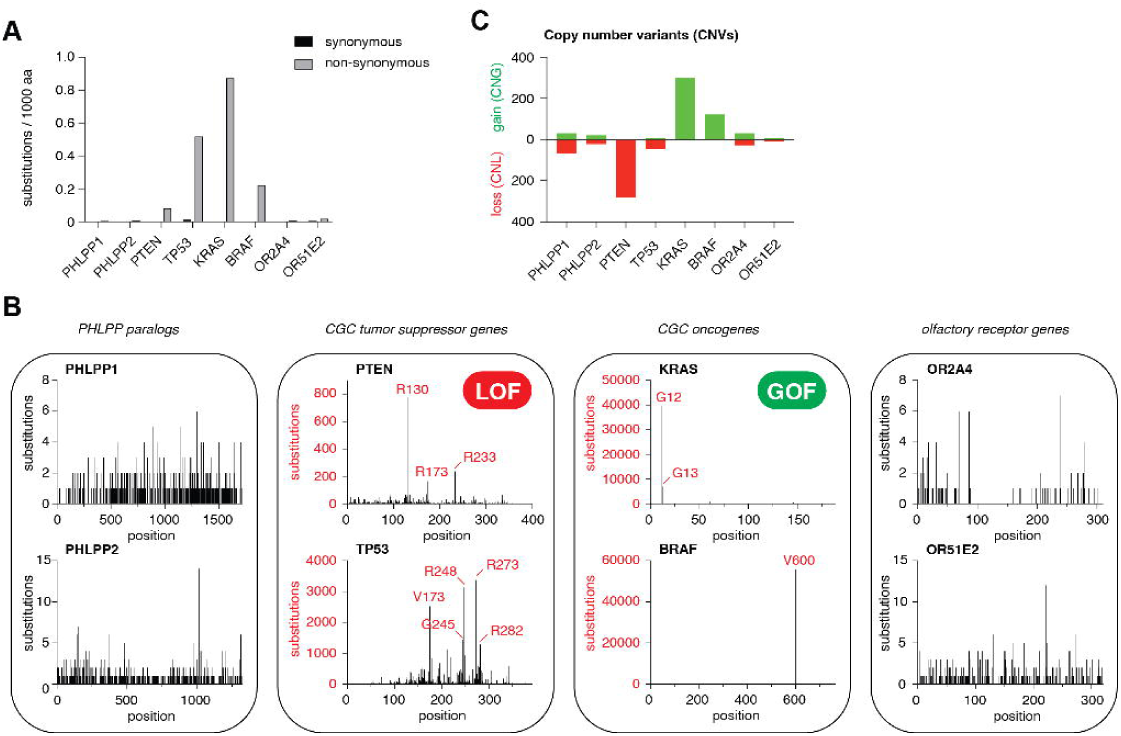
Cancer genomics does not support a role of PHLPP1 or PHLPP2 as tumor suppressors. A. Number of synonymous and non-synonymous substitutions per 1000 amino acids reported in COSMIC for: PHLPP1 and PHLPP2; the tumor suppressor genes PTEN and TP53; the oncogenes KRAS and BRAF; the olfactory receptor genes OR2A4 and OR51E2. B. Distribution of non-synonymous mutations within the coding sequence of the genes represented in A. Red: known hotspot mutations in Cancer Gene Census (CGC) genes. C. Copy number variants (CNV) for the genes represented in A. Green: copy number gains (CNG); red: copy number losses (CNL).

To assess whether PHLPP1 or PHLPP2 is an essential gene in cancer cells, we plotted the gene effect score for 1100 cancer cell lines reported in the Dependency Map (DepMap) Public 23Q4+Score (Chronos) CRISPR knockout screens (34–37), 710 cancer cell lines reported in the Achilles+DRIVE+Marcotte (DEMETER2) RNA interference (RNAi) knockdown screens (38–41), and 317 cancer cell lines reported in copy number effect-corrected (36) CRISPR knockout screens (Supplementary Figure 4A-B). Neither PHLPP1 nor PHLPP2 were found to be essential in any of these screens. These observations are consistent with the viability of single knockout mice for both PHLPP1 (42, 43) and PHLPP2 (14), as well as a *PHLPP1* ^-/-^ *PHLPP2* ^-/-^ double knockout mouse (14), none of which have been reported to exhibit a significantly higher propensity to develop tumors.

Finally, we mapped the results of genome-wide association studies (GWAS) for the genetic loci of *PHLPP1* and *PHLPP2*. An intron variant in the neighboring *BCL2* gene has been associated with prostate carcinoma in two GWAS studies with p-values < 5.0 x10^-8^ (44, 45), but no cancer associations have been reported for either *PHLPP1* or *PHLPP2* (Supplementary Figure 4C-D). Analysis of synthetic lethal screens of cancer cells also found no evidence for synthetic lethal interactions involving either PHLPP1 or PHLPP2 (46). In summary, there is scant evidence from disease genotypes, clinically reported variants, or GWAS studies to support that dysregulation of either PHLPP1 or PHLPP2 is associated with cancer.

### PHLPP2 exhibits a conserved arrangement of its regulatory domains

Both PHLPP1 and PHLPP2 comprise an N-terminal Ras-associated (RA) domain followed by a PH domain, LRR domain, and a C-terminal protein phosphatase 2C (PP2C) domain (Figure 5A). Rotary shadowing electron microscopy revealed particles with a regular horseshoe-shaped conformation (Figure 5B, inset) consistent with the AlphaFold2-predicted structure of PHLPP2 (Figure 5C-D). We next determined the structure of PHLPP2 by single particle cryo-electron microscopy (cryoEM). We obtained two distinct reconstructions from the sample (see Supplementary Figure 5 for sample micrographs, 2D class averages, and Fourier Shell Correlation curves). We interpret these as endpoints of a conformational continuum, which explains why the resolution of both maps is limited to 6 Å. The two conformations differ in the location of the PH domain, which appears to be very flexible and is the least resolved part of the structure. Furthermore, the LRR exhibits bending while the PP2C and RA domains show compaction, potentially changing the accessibility of the phosphatase domain. Nevertheless, the AlphaFold2 prediction of PHLPP2 fits well into the particle volumes for both conformations (Figure 5E-F).

**Figure 5.**
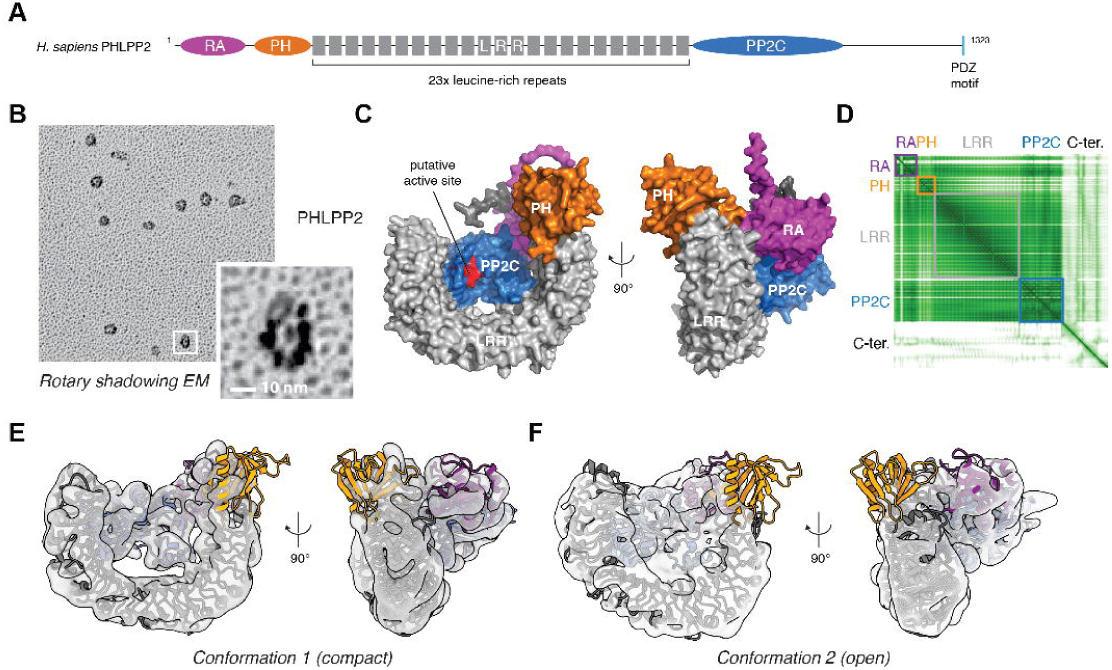
PHLPP exhibits a conserved arrangement of its regulatory domains. A. Domain architecture of human PHLPP2. (RA domain, magenta; PH domain, orange; LRR (grey); PP2C, blue; putative active site of PP2C domain, red). B. Rotary shadowing electron microscopy of recombinant PHLPP2. C. Top-ranked AlphaFold22 model of human PHLPP2. D. Pair alignment error (PAE) plot for AF2 prediction in E. PHLPP2 domain boundaries are indicated by colored boxes. E. CryoEM maps of human PHLPP2 (conformation 1) at 6 Å resolution with the AlphaFold2-predicted structure of PHLPP2 fit to the map. F. CryoEM maps of human PHLPP2 (conformation 2) at 6 Å resolution with the AlphaFold2-predicted structure of PHLPP2 fit to the map.

### Evolution and diversification of PHLPP genes

To provide clues to PHLPP function, we used phylogenetics to reconstruct the evolutionary history of the PHLPP protein family. The presence of PHLPP homologs in distantly related eukaryotes such as amoebozoans and parabasalids, suggests that the PHLPP family emerged early in eukaryotic evolution, presumably following the fusion of an LRR protein with a PP2C family phosphatase (Figure 6A-B). The ancestral gene encoded all residues known to be required for catalytic activity, including coordination of the M1 and M2 metal ions (Figure 6C). In the last common ancestor of opisthokonts (e. g., animals and fungi), around 1000 Mya, the ancestral LRR-PP2C gene gained a RA domain and was subsequently duplicated. Shortly thereafter, one copy of the gene gained a PH and a kinase domain giving rise to the PHLPP family, whereas the other paralog gained a class III adenylate cyclase (AC), resulting in the emergence of the Cyr1 family (Figure 6B). Retention of PHLPP and Cyr1 in unicellular holozoans (*P* = 4.56×10^-5^, Δ log-likelihood = 114.7, approximately-unbiased (AU) test for monophyly), the closest unicellular relatives of animals, indicates that this duplication was followed by reciprocal loss of Cyr1 in metazoa and PHLPP in fungi. The ancestral Cyr1 was likely a pseudophosphatase given the lack of catalytic residues, but it later acquired a dimerization domain in fungi (Figure 6A-C). This contrasts with PHLPP, which only lost its active site in metazoa. Indeed, PHLPP homologs in unicellular holozoans not only encode active site residues but in some cases include a C-terminal kinase domain (Figure 6A-C). Nonetheless, PHLPP was conserved throughout animal evolution before giving rise to PHLPP1 and PHLPP2 following a duplication event in a common ancestor of vertebrates. To better understand the impact of these evolutionary changes on PHLPP structure, we employed AlphaFold2 to predict the three-dimensional structures of representative proteins from across the LRR-PP2C phylogeny. The LRR-PP2C phosphatase of *Entamoeba* is structurally consistent with each of the evolutionarily descended derivatives, including vertebrate PHLPP1 and PHLPP2 and fungal Cyr1 (Figure 6D). Together these data suggest that PHLPP emerged through a complex history of neo- and sub-functionalization, based on a conserved LRR-PP2C chassis which repeatedly lost phosphatase activity, including at the origin of animals.

**Figure 6.**
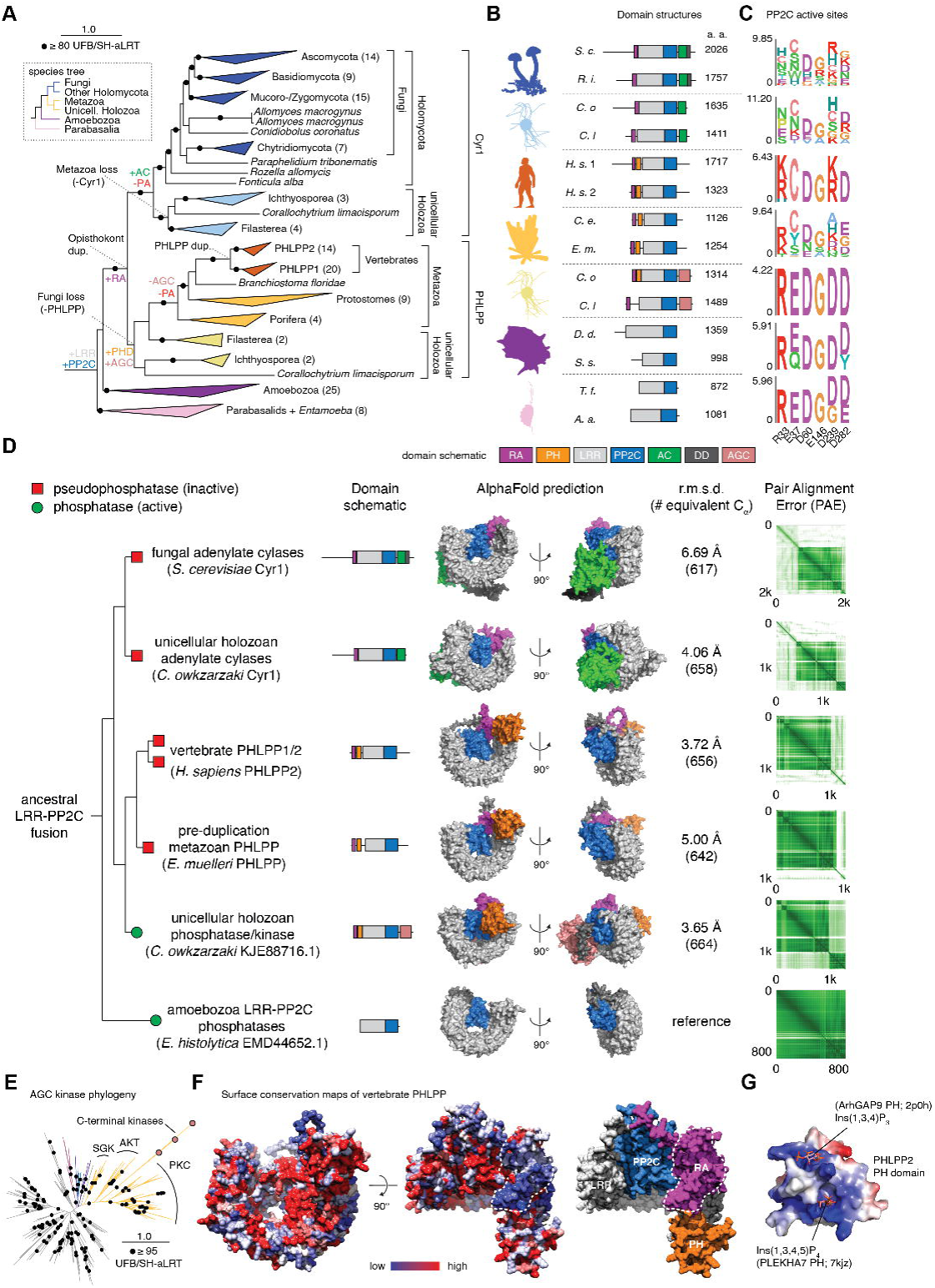
Evolution and diversification of PHLPP genes. A. Maximum-likelihood phylogeny of the LRR-PP2C protein family. Species and clade names have been added and the number of sequences within collapsed nodes are noted in parentheses. The gain and loss of genes, domains, and the PP2C active site (PA) has been mapped across the phylogeny based on Dollo parsimony. A species tree based on the taxa present in the phylogeny and current eukaryotic taxonomy (80) is shown for reference. The full phylogeny, domain annotations, and active site residues can be viewed at https://itol.embl.de/shared/9pitR7NesER8. B. Representative domain architectures of proteins from across the LRR-PP2C phylogeny. Taxonomic groups are noted with cartoons obtained from Phylopic.org and protein lengths are shown in amino acids (a. a.). *S. c.*, *Saccharomyces cerevisiae*; *R. i*.*, Rhizophagus irregularis; C. o., Capsaspora owczarzaki; C. l., Corallochytrium limacosporum; H. s. 1, Homo sapiens* PHLPP1; *H. s.* 2, *Homo sapiens* PHLPP2; *C. e.*, *Caenorhabditis elegans*; *E. m.*, *Ephydatia muelleri*; *D. d.*, *Dictyostelium discoideum*; *S. s.*, *Stenamoeba stenapodia*; *T. f.*, *Tritrichomonas foetus*; *A. a., Anaeramoeba ignava*. RA, Ras binding domain; AC, adenylate cyclase; PH, PH-domain; DD, dimerization domain; LRR, leucine rich repeat; AGC, AGC kinase domain; PP2C, PP2C domain. C. Sequence logos showing PP2C active site conservation. PPM1A residues are noted for reference. D. Domain architectures, AlphaFold2-predicted structures, and their associated confidence metrics (PAE plots). The root mean square deviation (r.m.s.d.) of the LRR-PP2C domains (over C_α_ atoms) of each structure from the ancestral LRR- PP2C phosphatase of *E. histolytica* is shown for comparison. Red squares: pseudophosphatase (inactive); green circles: phosphatase (active). E. Maximum-likelihood phylogeny of kinase domains. The C-terminal kinase domains of unicellular holozoan Cyr1 cluster with AGC kinases of the Akt and PKC families. The kinase domain phylogeny can be viewed at https://itol.embl.de/shared/9pitR7NesER8. F. Surface conservation maps of vertebrate PHLPP. Low conservation (blue) to high conservation (red). Dashed white line: surface of RA domain used to bind RAS in Cyr1. G. Electrostatic surface potential map of the PH domain of H. sapiens PHLPP2. Phosphoinositide headgroup ligands superimposed from experimentally determined structures of homologous ArhGAP9 and PLEKHA7 PH domains identified by FoldSeek (66).

The structure and function of Cyr1 provides a useful comparison when considering the functional diversification of PHLPP. The acquisition of an RA domain, which preceded gene duplication at the base of the opisthokont clade, apparently gave rise to a Ras-dependent functionality in fungal Cyr1. *Saccharomyces cerevisiae* Cyr1 adenylate cyclase (AC) activity is stimulated by GTP-bound RAS1 or RAS2 (47), homologs of human KRAS. AlphaFold2 prediction of the RAS1/2:Cyr1 interaction converges on five identical models of very high confidence (Supplementary Figure 6A) that are superimposable with experimentally-determined structures of KRAS in complex with RA domains from AF6 (48) and RGL1 (49) (Supplementary Figure 6B).

Reconstitution of yeast Cyr1 adenylate cyclase activity in the heterologous host *E. coli* implies that Cyr1 and RAS are necessary and sufficient for cAMP production (47). Since RAS binding to the RA domain of Cyr1 is not predicted to induce any conformational change in the RA domain, an allosteric mechanism can, most likely, be ruled out. Class III adenylate cyclases, of which Cyr1 is a member, depend on the homo-, hetero-, or pseudo-dimerization of two cyclase domains for cAMP production (50). Farnesylation of RAS is essential for Cyr1 activation (51), suggesting that RAS.GTP-dependent recruitment of Cyr1 to the plasma membrane may drive dimerization of the Cyr1 adenylate cyclase domain. AlphaFold2 prediction of a putative dimer of full-length Cyr1 converges on near-identical models (Supplementary Figure 6C) in which the C-terminal ∼400 amino acids exhibit a dimeric arrangement (Supplementary Figure 6D) superimposable with the AC domains of Cya (Supplementary Figure 6E), a bacterial adenylate cyclase, the structure of which has been experimentally determined by cryo-electron microscopy (52).

The C-terminal domain (CTD) predicted to dimerize Cyr1 has been reported to bind cyclase-associated protein (CAP), a multi-domain protein which links Cyr1 to the actin cytoskeleton (53). Biochemical studies have narrowed down the region of CAP essential for Cyr1 binding to the N-terminal 36 amino acids (54). AlphaFold2 prediction of the complex between the CTD of Cyr1 and CAP converges on 5 identical and high-confidence models in which the dimeric CTD binds the N-terminal helix of CAP in a hetero-tetrameric arrangement (Supplementary Figure 6F). This model therefore reconciles the mode and stoichiometry of binding of CAP to Cyr1. By combining the predicted models of interaction of Cyr1, RAS and CAP, a high-confidence prediction of a Cyr1-RAS-CAP hetero-hexamer (2:2:2) can be obtained (Supplementary Figure 6G).

Similar to Cyr1, acquisition of a PH domain in the PHLPP family appears to have coincided with C-terminal neo-functionalization through the emergence of a kinase domain. Curiously, phylogenetics suggest that this domain belongs to the AGC kinase family, which includes AKT and protein kinase C, another previously proposed target of PHLPP (Figure 6E) (55). Like the adenylate cyclase of Cyr1, this kinase domain was appended to the PHLPP C-terminus and may be positionally flexible, as indicated by AlphaFold2 predicted alignment error (PAE) plots (Figure 6D). The kinase domain was subsequently lost in the metazoan ancestor, concurrently with the phosphatase active site. The functional implications of this domain remain unclear, but could imply an ancestral connection between PHLPP and AGC kinases. Despite the apparent pseudogenization of PHLPP, surface conservation maps of vertebrate PHLPP indicate two regions of high conservation: one on the surface of the PH domain and one at the interface of the LRR and PP2C domains (Figure 6F). Importantly, whilst the RA domain has been retained in vertebrate PHLPP, its binding to RAS has not. In fact, the surface of the RA domain used by Cyr1 to bind RAS exhibits the highest sequence divergence of any region in PHLPP1 and PHLPP2 (Figure 6F). The acquisition of a PH domain may have substituted for RAS binding, although in the unicellular holozoan, *Corallichytrium limacisporum*, the PH domain appears to have been lost (Figure 6A), suggesting a period of functional redundancy. Electrostatic surface potential analysis of PHLPP2 indicates a basic surface overlapping with the binding sites of phosphoinositides in structural homologs of the PH domain (56, 57) (Figure 6G), suggesting a role for PHLPP at cellular membranes. Surface conservation within the LRR-PP2C cradle overlaps with an insertion between strands β7 and β8 of the PP2C domain that is known to be important for substrate binding in other PP2C phosphatases. These observations imply that vertebrate PHLPP may have lost phosphatase activity, but retained binding to the substrate of the ancestral protein phosphatase.

## Discussion

### PHLPP2 is a pseudophosphatase

We show that purified full-length PHLPP2 exhibits no detectable activity against either T308 or S473 of Akt in vitro, even at concentrations 500-fold higher than that found in the cell and at substrate concentrations >1500-fold higher. Hypothetically assuming that PHLPP2 is the active species, the maximal catalytic rate observed in the absence of okadaic acid corresponds to 0.1 dephosphorylation reactions per second, an order of magnitude lower than reported for the PP2C domain of PHLPP2 purified from insect cells (18). Together with our identification of contaminating PP2A in preparations of PHLPP2, this highlights the need for more stringent purification conditions than have previously been employed in order to make confident assessments of the biochemical activity of PHLPP proteins.

Phosphatases of the PPM family catalyze hydrolysis of phosphorylated serine or threonine residues by virtue of a bi-nuclear catalytic site in which two manganese ions are coordinated by a combination of aspartate side chains and ordered water molecules (27). A bridging water molecule coordinated by both Mn^2+^ ions is poised for nucleophilic attack on the substrate phosphate (26, 27) and the M2 metal ion determines the rate-limiting step in substrate hydrolysis (58). Mutations that disrupt metal binding or divalent cations other than Mn^2+^ or Fe^2+^ (including Zn^2+^) either abolish or strongly inhibit PPM1A activity (26, 59, 60). The observation of a single zinc ion in PHLPP2 with a complete coordination sphere suggests that PHLPP2 is incapable of canonical, binuclear-catalyzed phosphoester hydrolysis. Curiously, the carboxyl group of D822 in PHLPP2 occupies the same position as the phosphate group of a phosphorylated substrate peptide crystallized in complex with PPM1A (26), suggesting that loss of phosphatase activity may have coincided with a mutation that created a high-affinity zinc-binding site. Reports of pharmacologically active small molecule inhibitors of PHLPP should therefore be treated cautiously (15, 17). Since all known metal dependent phosphatases of the PPM family, including ancient PPM homologs in cyanobacteria, depend on two or three divalent cations in their active sites for catalysis (61), the lack of phosphatase activity of PHLPP2 and the related PPM family pseudophosphatase TAB1 (29) is expected. On this basis, in fact, both PHLPP1 and PHLPP2 have previously been classified as pseudophosphatases (62). Although not explicitly tested in this study, the predicted loss of activity in the last common ancestor of metazoans implies that PHLPP1 is also a pseudophosphatase. This raises the obvious question of what function PHLPP fulfils in the cell.

### Evidence does not support a role of PHLPP1 and PHLPP2 in cancer

Conceptually, a tumor suppressive function of PHLPP follows logically from its identification as an Akt phosphatase. However, the lack of detectable phosphatase activity in vitro does not support this role. Reports characterizing PHLPP as a tumor suppressor have relied heavily on correlations between PHLPP expression levels and Akt phosphorylation in cancer cell lines (12, 13). Causation was originally inferred from observations that reintroduction of PHLPP into a glioblastoma cell line resulted in suppression of tumor growth (12). Knockdown of PHLPP2 in cells also resulted in a significant increase in Akt S473 phosphorylation, which was interpreted to be a direct consequence of the loss of phosphatase activity (13). In the context of our findings, however, some of the previously reported effects on Akt phosphorylation may have been a pleiotropic effect of the ectopic over-expression of PHLPP. In particular, dominant negative effects on membrane signaling cannot be ruled out given the reported subcellular localization of PHLPP to the plasma membrane (63). However, a recent study revealed that adipose tissue-specific ablation of PHLPP2 in mice as well as siRNA-mediated knockdown of PHLPP2 in 3T3-L1 adipocytes did not lead to any significant differences in Akt S473 phosphorylation (64), suggesting that PHLPP2 may not be involved in Akt signaling, even indirectly.

Oncogenes and tumor suppressor genes have been carefully curated in the Cancer Gene Census, regularly updated by the COSMIC curation team (33). This list, which now numbers more than 700 human genes, requires indisputable mutation patterns in specific cancers reported in at least two independent studies from different groups. Consistent with the lack of somatic mutations in cancer, background mutation rates comparable to olfactory receptor genes, and no patterns of copy number loss in cancer, neither PHLPP1 nor PHLPP2 qualifies as a tumor suppressor. Taken together with the observation that PHLPP2 is a pseudophosphatase, the proposed role of PHLPP in cancer deserves further scrutiny, with an emphasis on its non-catalytic functions.

### PHLPP evolved from an ancestral phosphatase

The evolution of PHLPP from an ancestral phosphatase into two structurally homologous, but possibly functionally unrelated proteins is an example of neo- and sub-functionalization of a protein scaffold. The acquisition of a RA domain early in evolution implies that ancestral homologs of PHLPP were once able to bind RAS, but later lost this ability, supported by the absence of PHLPP in the proximal interactome of human RAS proteins (65). It seems highly probable that the RA domain was retained, despite loss of RAS binding, due to the intimate contacts it makes between the PP2C and LRR domains. The loss of RAS binding coincides with the acquisition of a PH domain, which has been bioinformatically classified as a ‘weak’ PIP_3_-binder, with apparent localization to the plasma membrane under some conditions (63). Surface conservation analysis indicates strong conservation of basic residues in the known binding pockets of PH domains that specifically recognize phosphoinositides. On balance, it seems highly likely that both fungal Cyr1 and metazoan PHLPP are membrane-resident proteins, regulated by RAS and phosphoinositides respectively. In principle, this should narrow the search for components of whatever biological pathway PHLPP is active in.

That pathway is unlikely to involve adenylate cyclase, since this was an adaptation in the paralogous gene family to PHLPP. Although CAP is highly conserved in vertebrates, including its N-terminus, there is no evidence of any structural homolog of the CTD of Cyr1 in animals (66). Previous studies have identified a number of PHLPP interaction partners by affinity capture mass spectrometry (AC-MS), including Na+/H+ exchanger regulatory factor 1 (NHERF1) (67) and the WD repeat-containing proteins 20 (WDR20) and 48 (WDR48; also known as ubiquitin-specific protease (USP)1-associated factor 1), USP12, and USP46 (68–74). A study in which USP1 was immunoprecipitated identified PHLPP1 and PHLPP2 as putative interactors (75), while a proximity biotin identification (BioID) screen of PHLPP1 also identified USP1, WDR20 and WDR48 (76). Although Akt has not been reported as an interaction partner of PHLPP1 or PHLPP2 in any of these high-throughput screens, USP12 and USP46 have been reported to attenuate Akt signaling by reducing the rate of proteasome-dependent PHLPP degradation (77). In light of our findings, however, this hypothesis may need revising.

Finally, it should be noted that PHLPP1 was originally identified and annotated as suprachiasmatic nucleus (SCN) circadian oscillatory protein (SCOP), on account of the circadian pattern of its gene expression in the SCN (78). SCOP was later proposed to play a role in the regulation of mitogen-activated protein kinase (MAPK) signaling and memory formation in the hippocampus (79). This data, in combination with the possible ancestral connection between PHLPP and AGC kinases, implied by the presence of a kinase domain in early PHLPP homologs, could suggest that PHLPP regulates kinases through non-catalytic mechanisms or may participate in vestigial non-functional kinase interactions.

In summary, purified PHLPP exhibits no detectable phosphatase activity. Public cancer repositories and genome-wide datasets provide no evidence for a role of either PHLPP1 or PHLPP2 in cancer. Phylogenetic analysis indicates that the ancestral gene that gave rise to PHLPP was a bona fide phosphatase, but that this activity was lost early in evolution. Although genetic ablation of either or both genes in mice is not lethal and results in no reported predisposition to cancer, an evolutionary pressure of some sort has retained both PHLPP1 and PHLPP2 in vertebrates. Efforts to determine the biological pathways that PHLPP participates in will undoubtedly require a focus on its tissue expression patterns, subcellular localization, and interactome. Many clues almost certainly lie in its evolution.

## Acknowledgements

We thank M. Brandstetter and the Electron Microscopy Facility of the Vienna Biocenter Core Facilities (VBCF) for help collecting rotary-shadowed EM images of PHLPP2. We acknowledge Diamond for access and support of the cryo-EM facilities at the U.K. national electron Bio-Imaging Centre (eBIC), proposal BI25222-3, as well as local contact Alistair Siebert for support. Proteomics analyses were performed by the Mass Spectrometry Facility at Max Perutz Labs using the VBCF instrument pool. We would like to thank C. Bock (Center for Molecular Medicine, Vienna and Medical University of Vienna) for help with mining cancer-associated public databases, genome-wide datasets, and gene essentiality screens. This work was supported by Austrian Science Fund (FWF) grants P33066, P36212, and W1261 as well as a Scientific and Technological Cooperation Grant (Austria-Bulgaria) of the OEAD (BG 05/2023) to T.A.L. and a Bulgarian National Science Fund grant to T.D. (KP-06-Austria/1). K.M.S. was supported by an Austrian Academy of Sciences (OEAW) DOC PhD Fellowship. DH and the Haselbach lab are supported by Boehringer Ingelheim.

## Author Contributions

T.H., K.M.S., and S.A. performed all in vitro biochemical experiments. V.M. and L.P. performed the immunoprecipitation experiments from mammalian cells. A.B. collected the ICP-MS data. S.E.A., N.K. and T.D. performed quantum mechanical modeling. K.M.S. and I.G. prepared cryo-electron microscopy grids. K.M.S., I.G. and D.H. collected electron micrographs and performed image processing. D.A. and M.H. designed and carried out the differential alkylation mass spectrometry experiments. N.I. and T.L. performed phylogenetic analyses. J.V. purified PP2A. L.T. purified Akt1^3P^. F.M.S. performed molecular dynamics simulations. B.Z., S.H., E.O., T.D., D.H., N.I., and T.A.L. funded the study. T.A.L. conceived and designed the study with specific input from E.O. with respect to the removal of contaminating PP2A.

## Materials and Methods

### Protein expression and purification

Recombinant human PHLPP2 and mutants thereof were expressed as N-terminal GST fusion proteins in baculovirus-infected *Spodoptera frugiperda* (Sf9) cells using a pFastBac Dual vector. Cells were harvested and lysed in buffer containing 50 mM Tris, pH 7.5, 150 mM KCl, 1 mM TCEP, 2 mM MgCl_2_, 125 U DENARASE (c-LEcta), and protease inhibitor cocktail (Sigma, P8849). After 1 h incubation at 4°C, the lysate was clarified by centrifugation at 39,000 g, 30 min, 4°C. The supernatant was incubated with glutathione Sepharose 4B resin for 2 h, 4°C. After washing with lysis buffer, PHLPP2 was cleaved off the beads with TEV protease overnight. The eluate from the beads was then applied to a 1 ml HiTrap Q column (Cytiva) and PHLPP2 was eluted with a linear salt gradient. Fractions containing PHLPP2 were pooled, concentrated to 0.5 ml and applied to a Superdex 200 30/300 column equilibrated in 20 mM Tris, pH 7.5, 150 mM KCl, 1 mM TCEP. The peak fractions containing PHLPP2 were pooled, concentrated, flash frozen in liquid nitrogen, and stored at −80°C. Akt1^3P^ was prepared according to Truebestein et al, 2021 (22).

### Electron microscopy

Purified, recombinant PHLPP2 samples were diluted to a final concentration of 50 μg/mL in spraying buffer containing 100 mM ammonium acetate and 30% (v/v) glycerol, pH 7.4. After dilution, the samples were sprayed onto freshly cleaved mica chips and immediately transferred into a BAL-TEC MED020 high-vacuum evaporator equipped with electron guns. While rotating, the samples were coated with 0.7 nm platinum at an angle of 4–5°, followed by 10 nm carbon at 90°. The obtained replicas were floated off from the mica chips and picked up on 400 mesh Cu/Pd grids. Grids were inspected in an FEI Morgagni 286D transmission electron microscope operated at 80 kV. Electron micrographs were acquired using an 11 megapixel Morada CCD camera from Olympus-SIS. Detailed methods for the preparation of samples for cryo-electron microscopy and image analysis can be found in the Supplementary Information.

### Malachite Green assay (MGA)

MGA assays were carried out according to the protocol of the Serine/Threonine Phosphatase Assay System (Promega). Serial dilutions of purified PHLPP2 or PP2A were incubated with 400 nM phospho-AL peptide (GATMKpTFCGT) or phospho-HM peptide (HFPQFpSYSAS) (GenScript) in the presence or absence of 12.5 nM okadaic acid, 1 h at 25°C. For PHLPP2 immunoprecipitated from HEK293 cells, serial dilutions (five times 1:3 and one 1:10) were prepared in a 96-well plate. The samples were split into two groups, with one group supplemented with okadaic acid to a final concentration of 12.5 nM. All samples were combined with phosphorylated Akt1 activation loop peptide to a final concentration of 0.4 mM. Phosphate production was measured at 620 nm in a TECAN Spark® spectrofluorometer and converted into reaction rates (mol product/min) using a calibration curve derived from phosphate standards.

### pNPP phosphatase assay

Serial dilutions of purified PHLPP2 or lambda phosphatase (purified in-house) were incubated with 10 mM pNPP in 20 mM Tris, pH 7.5, 150 mM KCl, 1 mM TCEP and 2 mM MnCl_2_, in the presence or absence of 12.5 nM okadaic acid, 1 h at 25°C. The conversion of pNPP to pNP was measured at 405 nm in a TECAN Spark spectrofluorometer.

## Notes

### Competing Interest Statement

The authors have declared no competing interest.

### Summary of Updates

A typo in the abstract stated "1 Mya" (million years ago), when it should have said "1000 Mya".

