## Supplementary Information for "PHLPP2 is a pseudophosphatase that lost activity in the metazoan ancestor"

[1]Max Perutz Labs, Vienna Biocenter Campus (VBC), Dr. Bohr-Gasse 9, 1030, Vienna, Austria.

[2]Medical University of Vienna, Center for Medical Biochemistry, Dr. Bohr-Gasse 9, 1030, Vienna, Austria.

[3]Vienna BioCenter PhD Program, a Doctoral School of the University of Vienna and the Medical University of Vienna, A-1030 Vienna, Austria.

[4]Research Institute of Molecular Pathology, Vienna BioCenter, Vienna, Austria.

[5]Institute of Optical Materials and Technologies "Acad. J. Malinowski", Bulgarian Academy of Sciences, 1113 Sofia, Bulgaria.

[6]University of Chemical Technology and Metallurgy, 8 St. Kliment Ohridski Blvd, 1756 Sofia, Bulgaria.

[7]University of Natural Resources and Life Sciences, Department of Chemistry, Institute of Analytical Chemistry, Muthgasse 18, 1190, Vienna, Austria.

[8]University of Vienna, Center for Molecular Biology, Department of Biochemistry and Cell Biology, Vienna, Austria.

[9]CeMM Research Center for Molecular Medicine of the Austrian Academy of Sciences, Vienna, Austria.

[10]Medical University of Vienna, Institute of Artificial Intelligence, Center for Medical Data Science, Vienna, Austria.

[11]Faculty of Chemistry and Pharmacy, Sofia University "St. Kliment Ohridski", 1164 Sofia, Bulgaria.

[12]Gregor Mendel Institute (GMI), Austrian Academy of Sciences, Vienna BioCenter (VBC), Vienna, Austria.

Contains:

Supplementary Methods
Supplementary Figures 1-6
Supplementary Table 1
Supplementary References

### Supplementary Methods

#### PP2A purification

2L of yeast cells (YJV1181; gal. prom. TAP-Pph21 endogenous, BY 4741) were grown to a density of $OD_{600nm}$=1 in 1% raffinose and TAP-Pph21 was expressed for 4 hours by addition of 1% galactose, yielding a final cell density of $OD_{600nm}$=2-3. Cells were harvested by filtration and frozen in liquid nitrogen. Cells were broken up using a Freezer Mill™ cryogenic grinder (SPEX) and resuspended in 20 ml ice-cold lysis buffer (50 mM MES pH 6.5, 150 mM NaCl, 1 mM EDTA, 1% Triton X-100, Complete protease inhibitor cocktail Roche). The lysate was pre-cleared by centrifugation (15' at 16 000 g) and incubated with 300 µl IgG Sephrose™ 6 Fast Flow beads (Cytiva 17096901) for 2h at 4°C. Beads were packed into a chromatographic column and washed 1x with 20 ml lysis buffer and 3x with 20ml cleavage buffer (50 mM MES pH6.5, 150 mM NaCl, 0.1% NP40). Beads were taken up in 800 µl cleavage buffer and Pph21 was cleaved by incubating the beads for 2h at 16°C with 500 U TEV protease (NEB P8112S). Eluted Pph21 was mixed with glycerol to a final conc of 50% and stored at -20°C.

#### Mass photometry

The mass distribution of purified PHLPP2 particles was measured on a Refeyn TwoMP mass photometer at a concentration of 25 nM. The mass of the respective peaks was estimated by fitting a Gaussian distribution to the histogram of particle masses.

#### Immunoblotting

After incubation with PHLPP2 under different conditions, the phosphorylation state of $Akt1^{3P}$ was monitored by Western blotting, using antibodies against Akt1 pT308 (Cell Signaling Technology #C31E5E) and pS473 (Cell Signaling Technology #193H12). Briefly, samples were separated by SDS-PAGE on a 12% tris-glycine gel and blotted onto nitrocellulose (0.4 µm pore size). The membrane was blocked for 1 h at room temperature with 5% BSA in 20 mM Tris, pH 7.5, 150 mM NaCl, 0.1% Tween 20. Membranes were incubated with primary antibodies at a ratio of 1:1000, overnight at 4°C, followed by washing and incubation with IRDye 800CW Goat anti-Rabbit IgG Secondary Antibody (LICOR, 926-32211) at a ratio of 1:15000, 1 h at room temperature.

After washing, the membranes were imaged using a LICOR Odyssey CLx fluorescence imager.

#### Size-exclusion chromatography coupled to inductively-coupled plasma mass spectrometry

Water was purified employing an Ultra Clear basic reverse osmosis system (SGWasseraufbereitung und RegenierstationGmbH, Barsbuettel, Germany) and subsequently subboiled (MLS, Leutkirch). Formic acid Suprapure® (purity 98-100%, item no. 111670), nitric acid (purity for analysis EMSURE® Reag. Ph Eur, ISO, item no. 100456), acetic acid Suprapure® (purity 100%. Item no. 100066) and ammonia solution (purity for analysis EMSURE® Reag. Ph Eur, ISO, item no. 1005432) were purchased from Merck (KGaA, Darmstadt,Germany). Nitric acid was double subboiled and stored in perfluoralkoxy-polymer bottles. Lyophilized bovine erythrocytes Cu/Zn-superoxide dismutase (Cu/Zn-SOD, item no. S7571) was purchased from Sigma-Aldrich Chemie GmbH (Vienna, Austria).

Samples were stored at -80 °C prior to measurement. All vials and materials which came into contact with the sample, blank and standard solutions were pre-cleaned via soaking for 24 h in 10% and 24 h in 1% nitric acid followed by flushing with distilled water and drying under a class 100 laminar flow clean bench. Measurements were performed under clean room conditions (class 10000). Between the sample runs 1% formic acid solution was injected to eliminate carry over, followed by one buffer blank for assessment of possible contamination.

A NexSAR Speciation Analysis Ready HPLC system from Perkin Elmer (Massachusetts, USA) controlled by Clarity 8.8 software was used for the size exclusion separation. The mobile phase consisted of 50 mmol/L ammonium acetate at pH 7 and the flow rate was set to 0.4 mL min$^{-1}$. The injection volume for injection of samples, blank solutions and standards onto the ACQUITY UPLC Protein BEH 450Å SEC column (Waters™) was 15 µL. For elemental detection, the system was combined with an NexION 2000 quadrupole ICP-MS from Perkin Elmer (Massachusetts, USA). The reaction cell was filled with oxygen to induce the formation of $^{32}S^{16}O^{+}$ and circumvent the $^{16}O_2^{+}$ interference on the most abundant sulfur isotope $^{32}S$. Zinc was detected as $^{66}Zn$. Bovine Cu/Zn-SOD was employed to determine the intensity ratio between the $^{66}Zn$ and the $^{32}S^{16}O$ signals. The stoichiometric zinc ratio was calculated from the number of sulfur atoms per protein. ICP-MS parameters and data evaluation were optimized according to (*1*). pH measurements were conducted with an Education Line EL20 pH-Meter with a micro electrode (inLab Micro, Mettler, Toledo, USA).

### Phosphatase inhibitor beads (PIBs)

PIBs were prepared according to (*2*).

### Molecular dynamics

The geometry of the predicted $Zn^{2+}$ binding site in PHLPP2 was determined using a two-step procedure involving explicit-solvent molecular dynamics (MD) simulation followed by quantum-mechanical optimization. For MD simulations of the full-length PHLPP2, the GROMACS simulation package v. 2023 (*3*), the all-atom Amber99SB-ILDN force field (*4*) and the TIP4P-D water model (*5*) were used. The initial configuration of the full-length PHLPP2 protein was obtained from an AlphaFold (*6*) prediction for PHLPP2, including manual placement of the zinc ion in the proposed active site. To prepare the system, an energy minimization, a 10-ns constant volume equilibration (NVT) and 10-ns constant pressure equilibration (NPT) were first performed without distance restraints using the parameters given below and reaching the final all-atom RMSD with respect to the AlphaFold structure of 0.13 nm. The system was simulated in a cubic water box (8 nm x 8 nm x 8 nm) with 0.15 M NaCl to mimic experimental conditions. We also studied the influence of $Zn^{2+}$ binding on the stability of the whole PHLPP2 structure. Due to inherent limitations, MD simulations cannot capture the quantum behavior of the metal-ion binding sites. Thus, distance restraints (k= 5000 kJ/mol/nm$^2$) between coordinating atoms and the zinc ion as well as between coordinating atoms themselves were included to stabilize the tetrahedral arrangement using the target distances from the X-ray structure of $\alpha$-klotho (*7*). The restrained simulations were run for a total length of 100 ns. A leap-frog algorithm was used for integration under periodic boundary conditions. In both energy minimization and production runs, neighbor-lists were updated every 20 steps, following a Verlet-scheme based grid-search approach. The bonds involving H atoms were constrained using LINCS (*8*). Temperature control was achieved using a V-rescale thermostat (*9*), with a relaxation time of 0.1 ps, while pressure was controlled using a Parrinello-Rahman barostat (*10*). Compressibility for the barostat was set to 4.5

x $10^{-5}$ and the relaxation time was 2 ps. Coupling was done separately for water and protein in all cases. A twin-range spherical cut-off (1.0 nm/1.2 nm) was used for van der Waals interactions, while electrostatics were treated using the Particle-Mesh Ewald (PME) approach with a real space cut-off of 1.2 nm, a 0.16-nm grid and cubic interpolation. During the restrained simulation, the all-atom RMSD with respect to the to the final structure of NPT equilibration reached a stable level of 0.2-0.25 nm after 10 ns (Supplementary Figure 2F), suggesting that the placement of the zinc ion in the M2 site does not affect the structural integrity of the rest of the PHLPP2 structure. Similarly, the RMSD of coordinating atoms relative to their positions in the initial structure for the restrained simulations remains under 0.05 nm throughout the simulation (Supplementary Figure 2H). Further quantum-mechanical optimization was performed starting with a structure from the pool with the lowest 5% in binding site RMSD with respect to the equilibrated unrestrained geometry.

### Quantum mechanical modeling (ONIOM)

Calculations were performed with the Gaussian 16 (Revision C.01) quantum chemistry software package (*11*). To reduce the computational time given the size of the system, we employed the ONIOM method (*12*) as implemented in the Gaussian 16 program. A two-layer QM/QM method that combines different QM methods is applied: a high-level of accuracy for the atoms three bonds away from the zinc ion and a low level for the rest of the systems – cut from the corresponding initial structures (crystal structure of a 1:1:1 FGF23-FGFR1c-α-Klotho Ternary Complex, PDB 5W21, and MD generated model) at approximately 12 Å in radius around the metal cation where the amino acid chains logically end with methyl groups. In all calculations for the α-Klotho model system, a DFT function (M062X, B3LYP, or ωB97XD) was adopted for the inner layer, while a semi-empirical method (PM3) was used for the treatment of the real system. Different basis sets (6-31G(d,p), 6-31+G(d,p) and 6-31+G(3d,p)) in conjunction with the selected DFT functions were assessed for an accurate prediction of structural parameters (Zn-S and Zn-O bond lengths). The sulfur atom of the cysteine was modeled in its deprotonated state, in accordance with the known interaction of sulfur with zinc, which has been theoretically predicted (*13*) and experimentally visualized in atomic resolution crystal structures. All the methods tested predicted Zn-S bond lengths in the range 2.32-2.40 Å, and the average lengths of the three Zn-O bonds were 1.96-2.02 Å. It was concluded that both oniom(ωB97XD/6-31G(d,p):PM3) and oniom(B3LYP/6-31+G(3d,p):PM3) methods are suitable for proper description of the geometry of the α-Klotho Zn center.

### Sample preparation for tandem mass spectrometry

Pellet of 1L Sf9 culture was lysed in 20 mL lysis buffer (2 mM benzamidine, 200 µL Protease inhibitor stock, 2mM MgCl2, 1 mM TCEP, 0.4 µL benzonase, 0.25% Chaps) for approximately 1 h at 4˚C in the cold room and then spun down (30 min, 18k rpm, 4˚C). The clarified supernatant was assessed for protein concentration using the Bradford assay, and two equal aliquots (1 mg/mL) were prepared. As a control, 0.5 µM okadaic acid was added to one of the samples. The two lysate samples (1 mL each) were then incubated with 20 µL of phosphatase inhibitor beads (PIBs) at 4°C for 1 hour. After incubation, the supernatant was carefully removed, and the PIBs were washed three times with 0.5 mL of Lysis buffer. Subsequently, the washed beads were incubated with 120 µL of 2% SDS for 1 hour at 65˚C and the eluates collected using Pierce Micro-Spin columns.

The samples were mixed with 5µL 100mM Tris HCl pH 8.5 and heated at 95°C for 5min. Then disulfide bonds were reduced with 4 µL of 250 mM dithiothreitol (DTT) for 30 min at room temperature before adding 4 µL of 500 mM iodoacetamide and incubating for 30 min at room temperature in the dark. The remaining iodoacetamide was quenched with 2 µL of 250 mM DTT for 10 min. Sera-MagSpeed Beads mix (GE Life Sciences, cat. nos. 45152105050350 and 65152105050350) was added to the sample at beads:protein ratio of 10:1 and mixed with ethanol to reach 50% final ethanol concentration. The tubes were kept in a thermomixer for 10min at 1000rpm for 24min to let protein bind to the beads. Then the supernatant was removed using a magnetic rack and the beads were washed with 180 µL 80% ethanol in water off the rack. And the wash procedure was repeated 3 more times. Afterwards, 30µL 50mM ammonium bicarbonate with 120 ng trypsin (Trypsin Gold, Promega) was added to the beads and incubated at 37°C 1000rpm overnight. The beads were settled on a magnetic rack and the supernatant was transferred to a new tube. The beads were rinsed with 30µL ammonium bicarbonate and sonicated for 30s and the supernatant was pooled to the supernatant from the previous step. Then the samples were centrifuged at 20000g for

1min and the supernatant was transferred to a new tube to remove any remaining beads. Trifluoroacetic acid was added to reach a final concentration of 0.5 %.

For the analysis of recombinant PHLPP2, ~5 µg protein (5 µL) was mixed with 15 µL 8M urea 50mM ammonium bicarbonate. Disulfide bonds were reduced with 0.8 µL of 250 mM dithiothreitol (DTT) for 30 min at room temperature before adding 0.8 µL of 500 mM iodoacetamide and incubating for 30 min at room temperature in the dark. The remaining iodoacetamide was quenched with 0.4 µL of 250 mM DTT for 10 min. 50mM ammonium bicarbonate was added to dilute the urea to 1M. The protein was digested with 160 ng trypsin (Trypsin Gold, Promega) at 37°C overnight. The digest was stopped by adding trifluoroacetic acid to reach a final concentration of 0.5 % and the peptides were desalted using C18 Stagetips (*14*).

For the differential alkylation experiment purified PHLPP2 was incubated in 5 mM iodoacetamide for 30 minutes in the dark alkylating accessible thiol groups. Unreacted iodoacetamide was removed by buffer exchange to 50 mM ammonium bicarbonate using Bio-Rad Micro Bio-Spin™ columns. Solid urea was added to the eluate to reach a concentration > 6 M urea. Subsequently disulfide bridges were reduced adding tris (2-carboxyethyl) phosphine (TCEP) to a final concentration of 1 mM and incubated for 30 minutes. Free thiols were then modified with methyl methanethiosulfonate (MMTS) at a concentration of 5 mM. After diluting to 1 M urea with 50 mM ammonium bicarbonate the sample was digested with trypsin at a protease:protein ratio of 1:30 at 37°C overnight. The digest was stopped by adding trifluoroacetic acid to reach a final concentration of 0.5 % and the peptides were desalted using C18 Stagetips.
As reference samples, the protein was denatured in 6.4M urea, reduced in 1mM TCEP and free thiols alkylated either with 5 mM iodoacetamide or 5 mM MMTS for 30 minutes in the dark prior digest and clean-up as described above.

#### Liquid chromatography- tandem mass spectrometry

Peptides were separated on an Ultimate 3000 RSLC nano-flow chromatography system (Thermo-Fisher), using a pre-column for sample loading (Acclaim PepMap C18, 2 cm × 0.1 mm, 5 µm, Thermo-Fisher), and a C18 analytical column (Acclaim PepMap C18, 50 cm × 0.75 mm, 2 µm, Thermo-Fisher), applying a segmented linear gradient from 2% to 35% and finally 80% solvent B (80 % acetonitrile, 0.1 % formic acid; solvent A 0.1 %

formic acid) at a flow rate of 230 nL/min over 60min . Eluting peptides were analyzed on an Exploris 480 Orbitrap mass spectrometer (Thermo Fisher), which was coupled to the column with a FAIMS pro ion-source (Thermo Fisher) using coated emitter tips (PepSep, MSWil).

For analysis of peptides obtained from PIB eluates, the mass spectrometer was operated in DIA mode with the FAIMS CV set to -45, the survey scans were obtained in a mass range of 350-1200 m/z, at a resolution of 60k at 200 m/z and a normalized AGC target at 300%. 31 MSMS spectra with variable isolation width between 13 and 24 m/z covering 349.5-1200.5 m/z range including 1 m/z windows overlap, were acquired in the HCD cell at 30% collision energy at a normalized AGC target of 1000% and a resolution of 30k. The max. injection time was set to auto.

For analysis of peptides obtained from purified, recombinant PHLPP2, the mass spectrometer was operated in DDA mode with two FAIMS compensation voltages (CV) set to -45 or -60 and 1 s cycle time per CV. The survey scans were obtained in a mass range of 350-1500 m/z, at a resolution of 60k at 200 m/z, and a normalized AGC target at 100%. The most intense ions were selected with an isolation width of 1.2 m/z, fragmented in the HCD cell at 28% collision energy, and the spectra recorded for max. 100 ms at a normalized AGC target of 100% and a resolution of 15k. Peptides with a charge of +2 to +6 were included for fragmentation, the peptide match feature was set to preferred, the exclude isotope feature was enabled, and selected precursors were dynamically excluded from repeated sampling for 20 seconds.

For the differential alkylation experiments, the Exploris 480 Orbitrap mass spectrometer was run without FAIMS. The survey scans were obtained in a mass range of 350-1500 m/z, at a resolution of 60k at 200 m/z, and a normalized AGC target at 300%. The 10 most intense ions were selected with an isolation width of 1.2 m/z, fragmented in the HCD cell at 30% collision energy, and the spectra recorded for max. 100 ms at a normalized AGC target of 200% and a resolution of 15k.

### MS data analysis

Raw data were processed using Spectronaut software (version 18.3, https://biognosys.com/software/spectronaut/) with the DirectDIA+ workflow or FragPipe (version 19.1) (*15*). The Uniprot Spodoptera_frugiperda proteome (version

2023.03), PHLP2_HUMAN protein sequence, as well as a database of most common contaminants were used. The searches were performed with full trypsin specificity and a maximum of 2 missed cleavages at a protein and peptide spectrum match false discovery rate of 1%. Carbamidomethylation of cysteine residues were set as fixed, oxidation of methionine and N-terminal acetylation as variable modifications. The cross-run normalization was turned off and all other settings were left as default. For the differential alkylation analysis, modification of cysteines with iodoacetamide or MMTS were defined as variable modifications.

Computational analysis was performed using Python and the in-house developed Python library MsReport (version 0.0.19) (*16*). Protein intensity less than 1000 was set to missing to remove the low-quality quantification values. LFQ protein intensities reported by SpectroNaut were log2-transformed and normalized across samples using the ModeNormalizer from MsReport. The missing normalized LFQ intensity values were imputed by drawing random values from a normal distribution after filtering out contaminants, proteins with less than 2 peptides and less than 1 quantified values in at least one group. Using MsReport, iBAQ intensities were calculated from the raw intensities reported by FragPipe.

The proteomics data have been deposited to the ProteomeXchange Consortium via the PRIDE partner repository (*17*) with the dataset identifiers PXD045891, PXD045978 and PXD052551.

#### Immunoprecipitation of PHLPP2

HEK293 cells (5x$10^6$) were transiently transfected with a plasmid encoding N-terminally V5-tagged PHLPP2 using Lipofectamine 3000 according to the manufacturer's protocol. Cells were harvested after two days, lysed with 3 repeated freeze/thaw cycles in liquid nitrogen, and then resuspended in 500 µL lysis buffer: 50 mM Tris pH 7.5, 150 mM KCl, 1 mM TCEP, 0.25% CHAPS, 2 mM $MgCl_2$, 2 mM benzamidine, supplemented with 0.01 µL DENARASE® and 5 µL protease inhibitor cocktail, (Sigma P8849).

The lysate was combined with 25 µL of V5-trap magnetic beads (Chomotek v5tma) pre-equilibrated in wash buffer (50 mM Tris pH 7.5, 150 mM KCl, 1 mM TCEP) and incubated for 3 h at 4°C on a rotary plate. A magnetic rack was used to pull down the beads and the supernatant (lysate) was removed. The beads were resuspended in wash buffer (beads). The wash step was repeated 5 times. Samples of the lysate, beads, and supernatant were taken after each wash step. All samples were subjected to a malachite green assay (Sigma-Aldrich MAK307) as described in the Methods section of the manuscript.

For western blotting, the samples were resolved by SDS-PAGE on a 12% Tris-Glycine gel. Transfer onto a 0.2 μm nitrocellulose membrane was performed at 120 V and 4 °C for 2.5 h. The membrane was incubated with 5 % BSA in TBST for 1 h while gently shaking. V5-Tag rabbit monoclonal antibody (Cell Signaling, #13202) and β-actin mouse monoclonal antibody (Cell Signaling, #3700) were diluted 1:1000 in 3 % BSA in TBST to a final volume of 5 mL and incubated with the membrane overnight at 4°C. The membrane was washed 3 times with TBST for 5 min each. Goat anti-rabbit secondary antibody (LI-COR, 926-32211) and goat anti-mouse secondary antibody (LI-COR, 926-68070) were diluted 1:15000 in 3 % BSA in TBST to a final volume of 5 mL and incubated with the membrane in the dark for 1 h at room temperature while gently shaking. The membrane was washed 3 times with TBST for 5 min each and imaged using a LI-COR Odyssey CLx fluorescence imager.

### AlphaFold2 structure prediction

The structures of PHLPP2 and Cyr1 homologs, as well as the Cyr1 dimer and complexes of Cyr1 with RAS1/RAS2 and CAP were predicted with AlphaFold2 multimer_v3 (*6*) installed on a local cluster. The top five models output in each case were compared to each other for convergence by calculating the r.m.s.d. over equivalent Ca atoms and evaluated according to their respective pair alignment error plots (PAE).

### Cryo-electron microscopy

*Grid preparation.* Open-hole cupper 200 mesh grids overlayed with R 2/2 carbon film (Quantifoil) were used for plunge freezing. Grids were glow-discharged with a BalTec SCD 005 sputter coater for 60 s at 25 mA using residual air. 4 μl of the sample at concentration of 0.1 mg/ml was applied onto the carbon side and front-side blotted using the Leica GP2 plunge freezer. The sample was blotted for 2 seconds including the integrated blot sensor.

*Data collection.* Initial grid screening was performed on a 200kV Glacios microscope (ThermoFisher) using a Falcon III detector (ThermoFisher). For high-resolution structure determination, data collection was performed at Diamond Light Source Krios I using a K3 detector (Gatan). Images were collected at super-resolution pixel size of 0.53 Å/pix with a cumulative fluence of 35 $e^-/Å^2$ in a defocus range from 1.7 to 2.5 μm in 0.2μm increments using EPU (ThermoFisher).

*Image processing.* Images were preprocessed using the cryosparc (*18*) live pipeline. In short, micrograph movies were aligned, fourier binned by a factor of two and the CTF parameters were determined. Particles were picked initially via blob picking. Clustering these results in 2D led to new templates that have been subsequently been used for particle selection. Particles underwent 2 rounds of 2D clustering and an ab-initio model was generated and refined. These proprocessed particles were transferred to relion 5 (*19*). 3D Clustering generated two classes with distinct features but only one was refining to subnanometer resolution. Refinement was performed using Blush regularisation (*20*).

*Modelling.* Into the map an alphafold (*6*) model was placed using UCSF ChimeraX (*21*). Models were adjusted and refined using MDFF in Namdinator (*22*).

### Phylogenetic and sequence analysis

To examine the evolutionary history of PHLPP, we generated a database of genome-predicted eukaryotic reference proteomes from UniProt ($n$ = 181, downloaded 25 January 2024) (*23*). This database was taxonomically balanced by selecting the best two proteomes per genus based on BUSCO (Benchmarking Universal Single Copy Orthologues) completeness (*24*). For metazoans, fungi, and embryophytes, more strict taxonomic criteria were set by selecting the best proteome per phyla for metazoans ($n$ = 17), the best proteome per class for fungi (max two per phylum, $n$ = 12), and the best proteome per order for embryophytes (max three per class, $n$ = 11). To identify PHLPP homologs, the database was searched using Diamond BLASTp v2.0.9 with human PHLPP1 (UniProt accession: O60346) as a query (query coverage ≥ 25%, $E < 10^{-5}$, sensitive mode) (*25*). The resulting hits were aligned using MAFFT v7.520 trimmed using trimAl v1.4.rev15 with a gap threshold of 50%, and a maximum-likelihood phylogeny was inferred with IQ-Tree v2.2.6 using the LG4M substitution model with topology support assessed using Shimodaira-Hasegawa approximate likelihood ratio tests (SH-aLRT, $n$ = 1,000) and ultra-fast bootstraps ($n$ = 1,000) (*26*–*30*). The phylogenies were inspected manually in FigTree v1.4 (http://tree.bio.ed.ac.uk/software/figtree/) and paralogs were excluded. The resulting homologs were then used for two additional iterative BLAST searches with stricter E-values of $10^{-25}$, the hits from which were analyzed phylogenetically and curated as described. To improve taxonomic sampling, we conducted an additional search using the EukProt V3 database and incorporated the resulting hits to produce a complete set of LRR-PP2C homologues from diverse eukaryotes (*31*).

Phylogenetic analyses were used to reconstruct the evolutionary history of PHLPP. To this end, the identified PHLPP homologues were aligned with MAFFT using the high accuracy L-Ins-i algorithm and the alignment was trimmed as before. A maximum likelihood phylogeny was inferred using the LG4M+F+R7 substitution model which was selected based on Bayesian information criteria using ModelFinder (*32*). Statistical support was inferred as before using SH-aLRT and the phylogeny was rooted based on recent eukaryotic species phylogenies (*33*). The monophyly of the unicellular holozoans was assessed using an approximately unbiased (AU) test implemented in IQ-Tree using a constrained phylogeny inferred under the same model as the unconstrained tree (*34*). The phylogenies were visualized and annotated using iTOL v6 (*35*).

Protein domain annotations were accomplished by mapping profile hidden Markov models (HMM) obtained from the Pfam-A database to protein sequences using HMMER v3.4 ($E < 10^{-3}$) (*36, 37*). However, to improve HMM sensitivity, the mapped domains were extracted, aligned using MAFFT, and used to produce new HMMs. This process was repeated iteratively three times until consistent domain annotations were obtained. Active site residue conservation was investigated using sequence logos generated with Skylign using weighted counts (*38*).

Lastly, to identify the origin of the unicellular holozoan kinase domains, the reference database was searched using Diamond BLASTp ($E < 10^{-5}$, query coverage ≥ 50%, sensitive mode, max 250 targets per query sequence) and the holozoan kinases as a query. The kinase domains of the resulting proteins were excised and subsequently aligned with the PHLPP kinases using MAFFT L-Ins-i. The alignment was then trimmed using trimAl with a gap threshold of 50% and a phylogeny was generated using IQ-Tree as described above, using the LG4M+I+R10 substitution model selected with ModelFinder. Proteins were annotated based on BLAST searches against SWISS-PROT (*39*).

### Supplementary Figure Legends

#### Supplementary Figure 1. PHLPP exhibits a conserved arrangement of its regulatory domains.

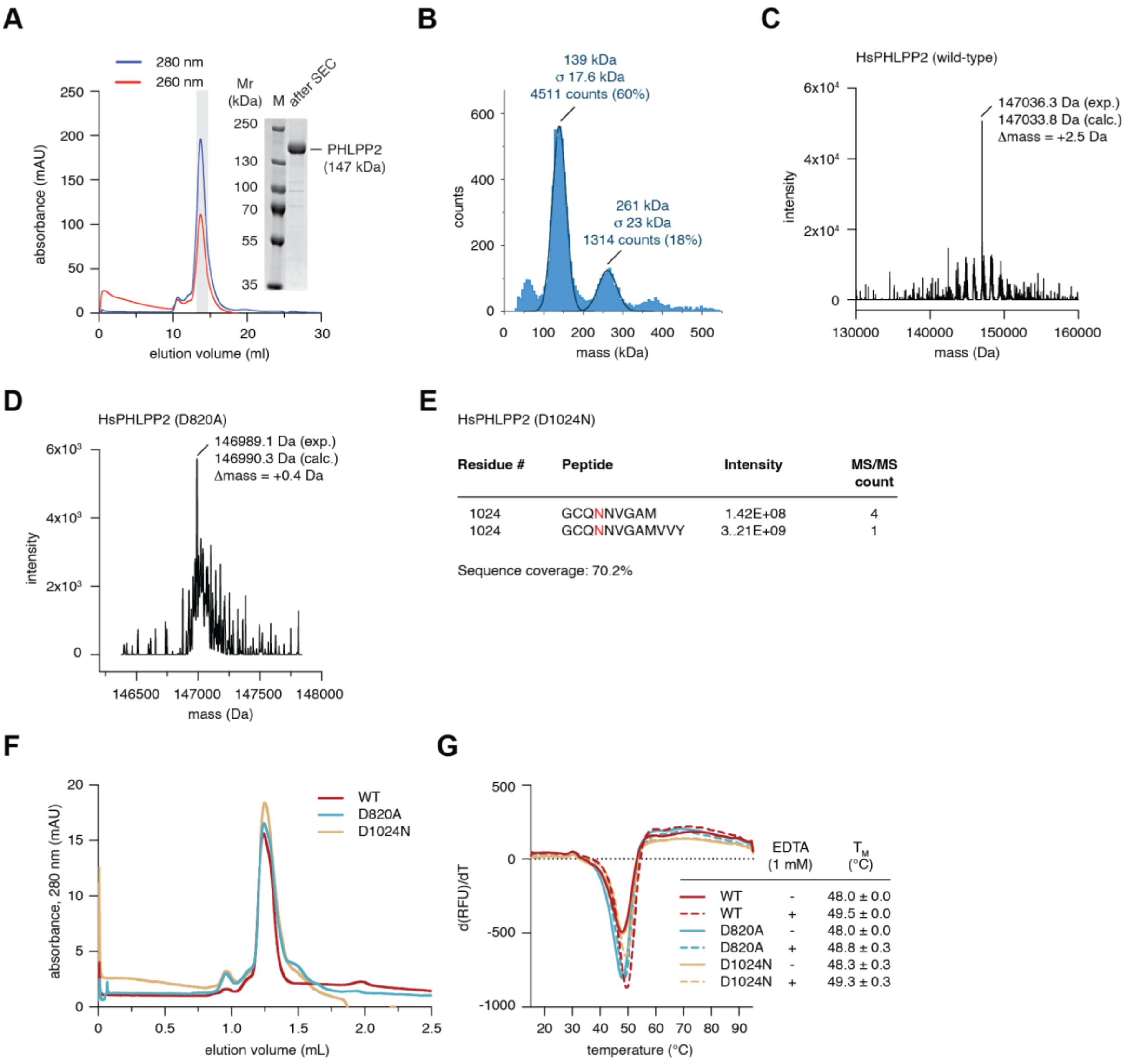


A. Size-exclusion chromatography of purified, recombinant human PHLPP2.

B. Mass photometry of recombinant PHLPP2.

C. Mass spectrum of recombinant human PHLPP2$^{WT}$.

D. Mass spectrum of recombinant human PHLPP2$^{D820A}$.

E. Peptide fingerprinting of recombinant human PHLPP2$^{D1024N}$. The -1 Da mass difference of the mutant was detected by tandem mass spectrometry analysis of chymotryptic peptides.

F. Analytical size exclusion chromatography of wild-type PHLPP2, PHLPP2$^{D820A}$ and PHLPP2$^{D1024N}$.

G. Thermal stability measurements of wild-type PHLPP2, PHLPP2$^{D820A}$ and PHLPP2$^{D1024N}$ in the presence and absence of 1 mM EDTA.

### Supplementary Figure 2. PHLPP2 exhibits no detectable activity against Akt.

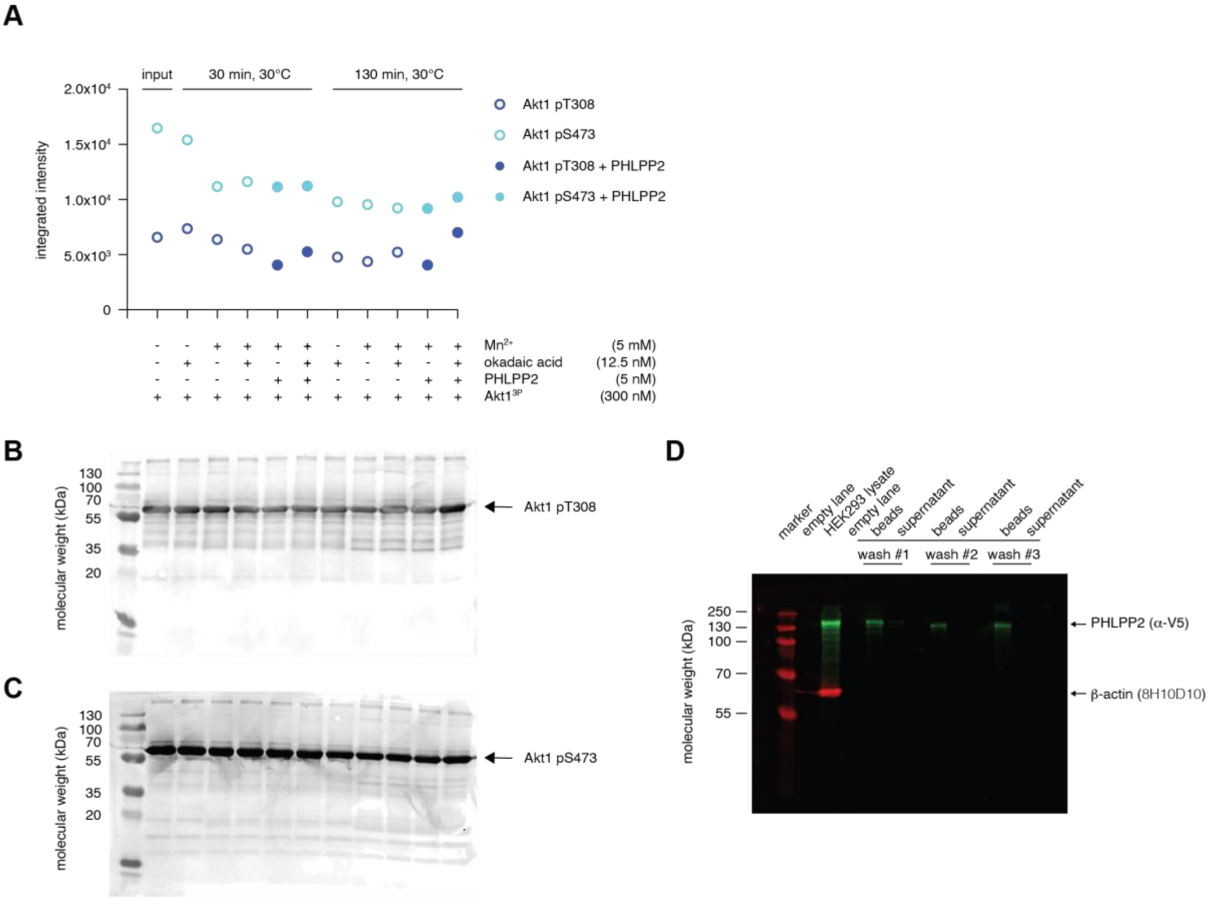


**E**

| Leading protein | Gene name | Protein name | Organism | Molecular weight (kDa) | Total # unique peptides | Sequence coverage | Imputed LFQ intensity [log2] -OA | +OA (ctrl) | Enrichment ratio [log2] |
|---|---|---|---|---|---|---|---|---|---|
| A0A2H1WUX1 | SFRICE_008486 | 60S ribosomal protein L24 | *Spodoptera frugiperda* | 17.39 | 2 | 13.50 | 25.37 | 15.00 | 10.37 |
| J9Z496 | | PP2A C-subunit | | 35.44 | 23 | 68.30 | 25.69 | 15.46 | 10.23 |
| A0A2H1W5F9 | SFRICE_005114 | Abl tyrosine kinase | | 63.75 | 2 | 4.80 | 20.68 | 10.74 | 9.94 |
| A0A2H1X028 | SFRICE_014915 | PP2A A-subunit | | 65.27 | 39 | 68.00 | 27.03 | 17.94 | 9.08 |
| A0A2H1WQP4 | SFRICE_007634 | PP2A 56 kDa γ-like subunit* | | 9.37 | 2 | 31.80 | 17.65 | 9.20 | 8.45 |
| A0A2H1VFZ6 | SFRICE_004402 | Protein Phosphatase 4 C-subunit | | 35.53 | 10 | 27.70 | 23.37 | 15.53 | 7.84 |
| A0A2H1WYQ1 | SFRICE_025240 | PP2A 56 kDa ε regulatory subunit | | 41.67 | 15 | 37.50 | 20.99 | 13.68 | 7.31 |
| A0A2H1W1C4 | SFRICE_000120 | uncharacterized protein | | 20.10 | 3 | 17.70 | 16.82 | 9.94 | 6.88 |
| A0A2H1V7H1 | SFRICE_001858 | Striatin-interacting protein (STRIPAK) | | 141.61 | 22 | 19.10 | 20.80 | 14.26 | 6.53 |
| A0A2H1VCR1 | SFRICE_002609 | Striatin-3 (PP2A B-subunit) | | 75.12 | 30 | 54.30 | 22.48 | 16.20 | 6.28 |
| A0A2H1V3Y5 | SFRICE_015530 | PP2A 55 kDa regulatory subunit B | | 56.78 | 24 | 45.10 | 23.73 | 17.53 | 6.20 |
| A0A2H1WL05 | SFRICE_022984 | MOB kinase activator-like 4 | | 25.51 | 9 | 36.70 | 22.69 | 16.74 | 5.95 |
| A0A2H1VWP9 | SFRICE_003851 | Protein phosphatase 6 regulatory subunit 3-A | | 91.52 | 17 | 20.10 | 23.07 | 17.66 | 5.42 |
| A0A2H1WE70 | SFRICE_008170 | Protein phosphatase 4 regulatory subunit 3 | | 90.35 | 23 | 31.30 | 18.20 | 13.49 | 4.71 |
| A0A2H1V4G9 | SFRICE_023010 | Myelin P2 protein | | 21.00 | 2 | 10.60 | 18.11 | 13.77 | 4.34 |
| A0A2H1X0P5 | SFRICE_029750 | Protein phosphatase 6 C-subunit | | 34.86 | 13 | 37.60 | 22.00 | 17.74 | 4.26 |
| A0A2H1W001 | SFRICE_003894 | Protein phosphatase 4 regulatory subunit 2 | | 54.77 | 20 | 46.30 | 19.80 | 15.59 | 4.21 |
| A0A2H1X3N4 | SFRICE_031227 | Microtubule-associated protein RP/EB 1 | | 41.94 | 2 | 4.80 | 15.15 | 10.99 | 4.16 |
| Q6ZVD8 | Q6ZVD8 | PHLPP2 (heterologous over-expression) | *Homo sapiens* | 146.66 | 19 | 17.10 | 19.16 | 19.68 | -0.53 |

A. Akt1$^{3P}$ dephosphorylation by purified PHLPP2, according to the assay conditions of Gao et al (*40*). Quantification of western blots shown in panels B and C.

B. Western blot of Akt1 pT308.

C. Western blot of Akt1 pS473.

D. Western blot of samples analyzed for phosphatase activity in Figure 4H. Immunoblotting was carried out with mouse anti-β-actin antibody and a rabbit anti-V5 antibody followed by detection with AlexaFluor-647 anti-mouse IgG and AlexaFluor-488 conjugated anti-rabbit IgG secondary antibodies respectively.

E. Tandem mass spectrometry analysis of proteins bound to microcystin-LR-conjugated beads after incubation with cell lysates of Sf9 cells heterologously over-expressing PHLPP2. Red: protein phosphatases and protein-phosphatase-associated regulatory proteins.

Supplementary Figure 3. PHLPP2 is a zinc-binding protein.

A

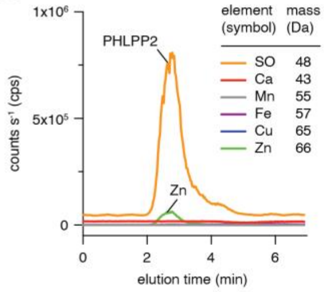


B

| Sample | $^{32}$S peak area (counts) | $^{65}$Cu peak area (counts) | $^{66}$Zn peak area (counts) | S/Cu ratio | S/Zn ratio | stoichiometry (S:Cu:Zn) |
|---|---|---|---|---|---|---|
| **SOD**[1] (calibrant) | 1324513 | 2253245 | 1854913 | 0.5878 | 0.7141 | 4:1:1 |
| | 1321345 | 2249440 | 1826849 | 0.5874 | 0.7233 | |
| | 1359483 | 2313953 | 1880422 | 0.5875 | 0.7230 | |
| Average | | | | 0.5876 | 0.7201 | |
| RSD (n=3) | | | | <1% | <1% | |
| **PHLPP2**[2] (replicate #1) | 50032296 | 448795 | 3674225 | 111.4815 | 13.6171 | 67 : 0.09 : 0.88 |
| | 48727540 | 415543 | 4204563 | 117.2623 | 11.5920 | 67 : 0.08 : 1.04 |
| | 18592408 | 111698 | 1685312 | 166.4532 | 11.0320 | 67 : 0.06 : 1.09 |
| Average | | | | | | 67 : 0.08 : 1.01 |
| RSD (n=3) | Total combined uncertainty of the measured stoichiometry: 11% | | | | | |
| **PHLPP2**[2] (replicate #2) | 38036679 | - | 2752163 | - | 13.8207 | 67 : - : 1.09 |
| | 23377197 | - | 1600115 | - | 14.6097 | 67 : - : 1.03 |
| | 55531332 | - | 4939412 | - | 11.2425 | 67 : - : 1.35 |
| Average | | | | | | 67 : - : 1.16 |
| RSD (n=3) | Total combined uncertainty of the measured stoichiometry: 14% | | | | | |
| **PHLPP2**[2] (replicate #3) | 350283355 | - | 35552871 | - | 9.8525 | 67 : - : 1.25 |
| | 403417464 | - | 42803314 | - | 9.4249 | 67 : - : 1.31 |
| Average | | | | | | 67 : - : 1.28 |
| RSD (n=2) | Total combined uncertainty of the measured stoichiometry: 10% | | | | | |

[1]Bovine SOD1; 3 Cys + 1 Met = total 4S.
[2]Human PHLPP2; 40 Cys + 27 Met = total 67S.

C

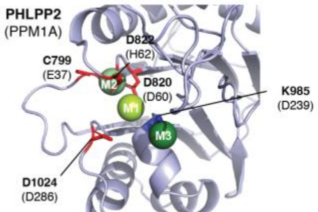


D

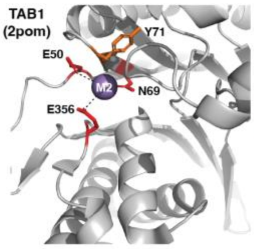


F

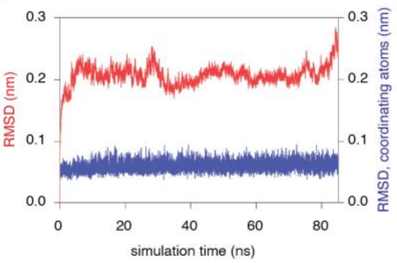


G

| ONIOM validation (α-klotho) | Bond: S - $Zn^{2+}$ (Å) | O - $Zn^{2+}$ (Å) |
|---|---|---|
| experimental structure (5W21) | 2.30 | 1.99 |
| B3LYP/6-31G(d,p):PM3 | 2.34 | 1.97 |
| B3LYP/6-31+G(d,p):PM3 | 2.36 | 2.02 |
| B3LYP/6-31+G(3d,p):PM3 | 2.34 | 2.01 |
| MO62X/6-31G(d,p):PM3 | 2.39 | 1.99 |
| MO62X/6-31+G(d,p):PM3 | 2.39 | 2.00 |
| MO62X/6-31+G(3d,p):PM3 | 2.40 | 2.01 |
| ωB97XD/6-31G(d,p):PM3 | 2.33 | 2.96 |
| ωB97XD/6-31+G(d,p):PM3 | 2.32 | 2.01 |

E

| Residue # (CYS) | Structure prediction | coverage (Y/N) | MMTS/IAA ratio | solvent accessible / inaccessible? |
|---|---|---|---|---|
| 133 | solvent accessible (RA) | Y | 0.23 | accessible |
| 143 | solvent accessible loop (RA) | Y | 0.01 - 0.02 | accessible |
| 177 | PH-LRR interface | Y | n.d. | n.d. |
| 180 | PH-LRR interface | Y | n.d. | n.d. |
| 189 | solvent accessible (PH) | Y | 0 | accessible |
| 258 | solvent accessible (RA-LRR) | Y | 0.27 | accessible |
| 402 | solvent accessible (LRR) | Y | 0.06 | accessible |
| 460 | solvent accessible (LRR) | Y | n.d. | n.d. |
| 467 | core (LRR) | Y | 25.46 - 41.28 | inaccessible |
| 516 | solvent accessible (LRR) | Y | 0 | accessible |
| 522 | core (LRR) | Y | 3.67 | partially accessible |
| 690 | core (LRR) | Y | 17.94 - 34.16 | inaccessible |
| 799 | solvent accessible (PP2C) | Y | 10.62 | inaccessible |
| 833 | solvent accessible (PP2C) | Y | 0.05 - 0.07 | accessible |
| 878 | core (PP2C) | Y | n.d. | n.d. |
| 901 | core (PP2C) | Y | n.d. | n.d. |
| 906 | core (PP2C) | Y | n.d. | n.d. |
| 948 | solvent accessible (PP2C) | Y | 6.5 | |
| 949 | solvent accessible (PP2C) | Y | 9.21 | |
| 955 | solvent accessible (PP2C) | Y | 0.47 - 3.38 | partially accessible |
| 1118 | solvent accessible (C-ter.) | Y | 2.83 - 2.87 | partially accessible |
| 1173 | solvent accessible (C-ter.) | Y | 0.13 - 0.22 | accessible |
| 1174 | solvent accessible (C-ter.) | Y | 0.13 - 0.22 | accessible |
| 1193 | solvent accessible (C-ter.) | Y | 0.14 - 1.45 | accessible |
| 1202 | solvent accessible (C-ter.) | Y | 0.04 - 0.10 | accessible |
| 1234 | solvent accessible (C-ter.) | Y | 0.00 - 0.01 | accessible |
| 9 | solvent accessible (N-ter.) | N | - | - |
| 32 | core (RA) | N | - | - |
| 67 | core (RA) | N | - | - |
| 77 | core (RA) | N | - | - |
| 319 | core (LRR) | N | - | - |
| 331 | solvent accessible (LRR) | N | - | - |
| 351 | solvent accessible (LRR) | N | - | - |
| 613 | solvent accessible (LRR) | N | - | - |
| 636 | core (LRR) | N | - | - |
| 722 | LRR-PP2C interface | N | - | - |
| 1014 | core (PP2C) | N | - | - |
| 1022 | core (PP2C) | N | - | - |
| 1040 | solvent accessible (C-ter.) | N | - | - |
| 1042 | solvent accessible (C-ter.) | N | - | - |

n.d. = not determined

H

| Residue | Bond | Molecular Dynamics (Å) | Quantum Mechanics: B3LYP/ 6-31+G(3,d,p):PM3 (Å) | ωB97XD/ 6-31G(d,p):PM3 (Å) |
|---|---|---|---|---|
| C799 | S - $Zn^{2+}$ | 3.0 | 2.34 | 2.33 |
| D820 | O - $Zn^{2+}$ | 1.8 | 2.04 | 1.96 |
| D822 | O - $Zn^{2+}$ | 1.9 | 2.03 | 1.97 |
| D1024 | O - $Zn^{2+}$ | 1.8 | 1.99 | 1.94 |

I

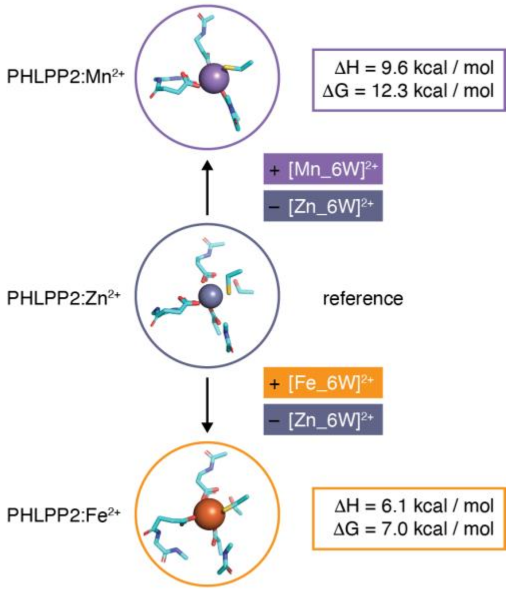

A. ICP-MS chromatograms for recombinant human PHLPP2, including the sulfur signal from the protein.

B. Calibration of PHLPP2:$Zn^{2+}$ stoichiometry for three biological replicates, each with duplicate or triplicate technical replicates.

C. Metal ion positions of PPM1A (PDB ID: 6b67) superimposed on AF2 model of PHLPP2 PP2C domain.

D. Experimentally determined structure of TAB1 pseudophosphatase, with one metal ion bound in the M2 position.

E. Tandem mass spectrometry analysis of differentially alkylated tryptic peptides from PHLPP2$^{WT}$ under native and denaturing conditions.

F. Cartesian root-mean-square deviation (RMSD) from the initial equilibrated structure over either all PHLPP2 atoms (red) or distance restrained groups (blue) as a function of time in MD simulations.

G. Validation of the ONIOM method for the experimentally determined C1D3 zinc finger of $\alpha$-klotho (*7*) (PDB ID 5W21). Comparison of bond lengths optimized by three quantum mechanical methods, each with different basis sets, to the experimentally determined bond lengths.

H. Comparison of interatomic distances between $Zn^{2+}$ and coordinating atoms in the MS2 binding site (carboxy atoms in aspartic acid or sulfur in cysteine) in the final MD structure and the final QM-optimized structure.

I. B3LYP/6-31+G(3d,p) fully optimized structures of metal binding sites of PHLPP2 comprising ligands from the first and second coordination layer, and enthalpies and Gibbs energies for $[M(H_2O)_6]^{2+}$ + $[PHLPP2\text{-}Zn]^{2-} \rightarrow [PHLPP2\text{-}M]^{2-}$ + $[Zn(H_2O)_6]^{2+}$ (M = Mn or Fe). Solvent accessible metal binding sites characterized with dielectric constant of 29 are considered. Calculation methodology according to (*41*).

Supplementary Figure 4. Cancer genomics does not support a role for PHLPP1 or PHLPP2 as tumor suppressors.

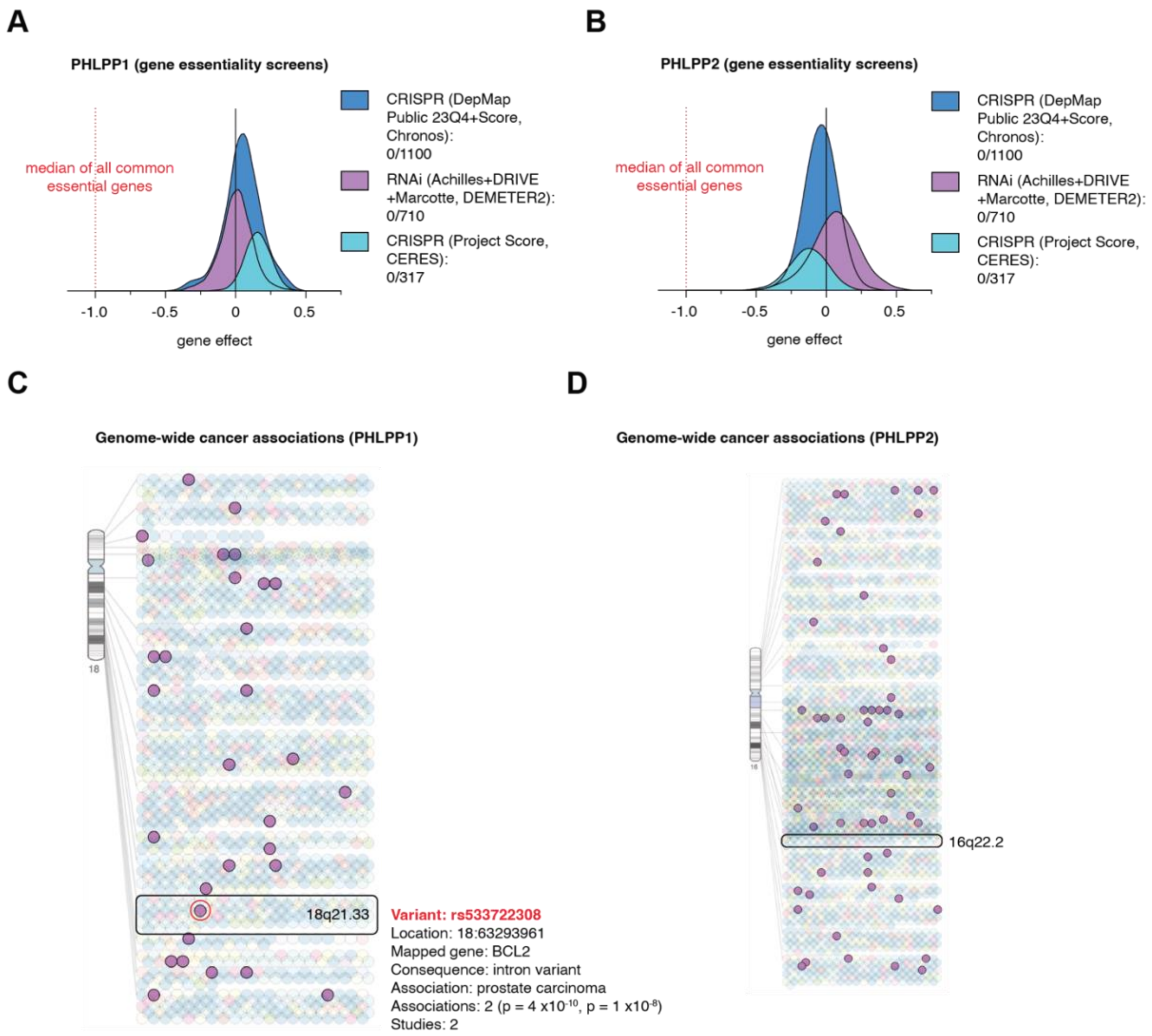


A. PHLPP1 essentiality determined from CRISPR knockout (blue) and RNAi (purple) screens for 1100 and 710 lines respectively. Copy number-corrected CRISPR knockout (CERES) gene effect scores for 317 cancer cell lines (cyan).

B. PHLPP2 essentiality determined from CRISPR knockout (blue) and RNAi (purple) screens for 1100 and 710 lines respectively. Copy number-corrected CRISPR knockout (CERES) gene effect scores for 317 cancer cell lines (cyan).

C. Genome-wide cancer associations in the chromosomal locus 18q21.33 containing the PHLPP1 gene. Red: single cancer-associated variant with a p-value < 5.0 x$10^{-8}$, which maps to an intron of the BCL2 gene.

D. Genome-wide cancer associations in the chromosomal locus 16q22.2 containing the PHLPP2 gene.

Supplementary Figure 5. Cryo-electron microscopy structure of PHLPP2.

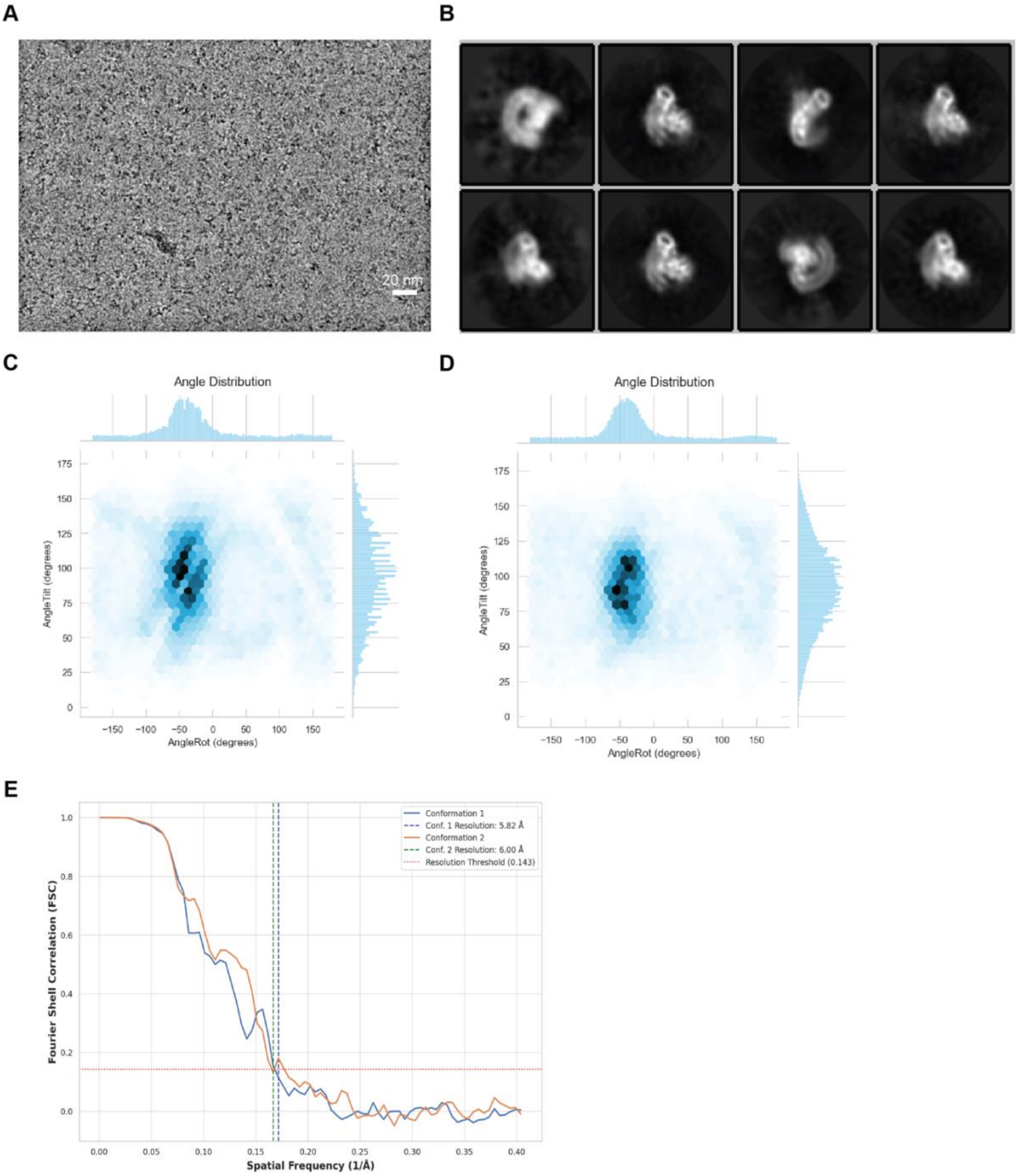


A. Representative micrograph.

B. Selected class averages representing the different viewing directions in the dataset.

C. Angle distribution plot of conformation 1.

D. Angle distribution plot of conformation 2.

E. Fourier shell correlation curves for each conformation.

Supplementary Figure 6. Evolution and diversification of PHLPP genes.

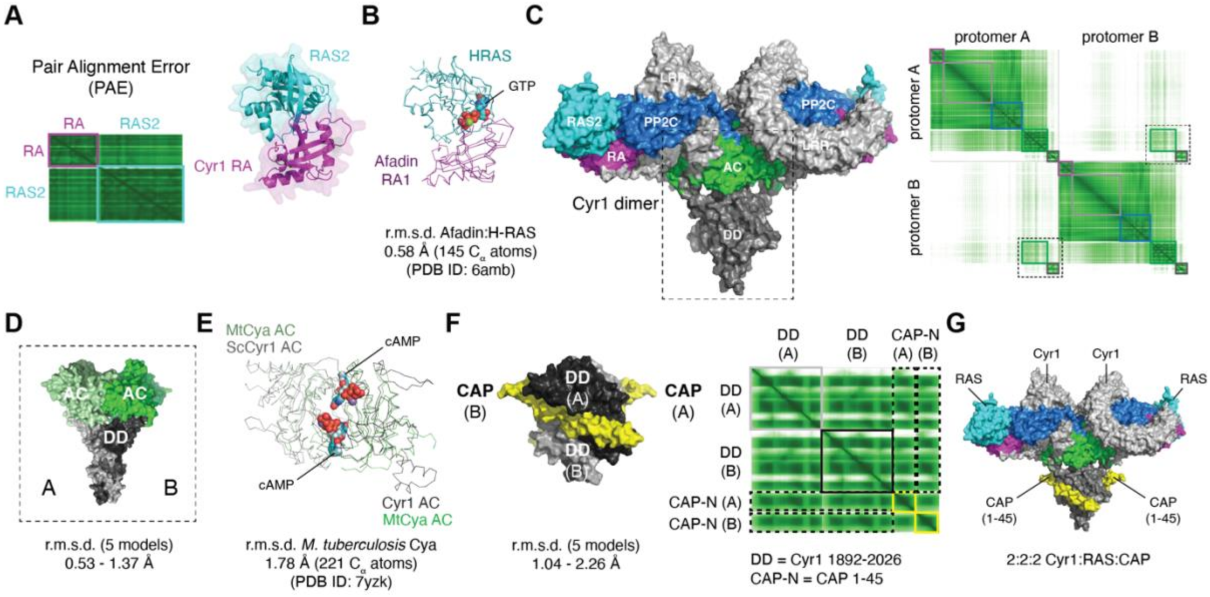


A. Structure prediction and associated PAE plot for the complex between the RA domain of *S. cerevisiae* Cyr1 (magenta) and RAS2 (cyan).

B. Superposition of the predicted Cyr1 RA:RAS2 structure (magenta/cyan) with the experimentally determined structure of the Afadin RA domain with GTP-bound H-RAS (purple/teal) (6amb). R.m.s.d. determined over 145 equivalent $C_\alpha$ atoms.

C. Structure prediction and associated PAE plot for dimeric *S. cerevisiae* Cyr1. Dashed rectangle: dimerization of the AC and CTD domains of Cyr1.

D. Structure prediction and associated PAE plot for dimeric AC-CTD domain of *S. cerevisiae* Cyr1. Agreement of 5 independent models shown by the r.m.s.d. over all $C_\alpha$ atoms.

E. Superposition of the predicted Cyr1 AC-CTD dimer (black/grey) with the experimentally determined structure of the AC domain of *M. tuberculosis* Cya (dark green/light green) (7yzk). R.m.s.d. determined over 221 equivalent $C_\alpha$ atoms.

F. Structure prediction and associated PAE plot for the dimeric CTD domain of *S. cerevisiae* Cyr1 (black/grey) in complex with two molecules of CAP (amino acids 1-45; yellow). Agreement of 5 independent models shown by the r.m.s.d. over all $C_\alpha$ atoms.

G. Structure prediction of the Cyr1:RAS2:CAP heterohexamer (2:2:2).

Supplementary Table 1. Cryo-EM data collection, refinement statistics

| | PHLPP conf1 (EMDB-51182) | PHLPP conf2 (EMDB-51183) |
|---|---|---|
| **Data collection and processing** | | |
| Magnification | 130 k | |
| Voltage (kV) | 300 | |
| Electron exposure (e–/Å$^2$) | 35 | |
| Defocus range (μm) | 1.7-2.5 | |
| Pixel size (Å) | 0.55 (super resolution) | |
| Symmetry imposed | C1 | |
| Initial particle images (no.) | 958089 | |
| Final particle images (no.) | 118375 | 355036 |
| Map resolution (Å) | 6.0 | 5.8 |
| FSC threshold 0.143 | | |
